# Tracking the trajectory of individual conjugative plasmids in bacterial communities using CRISPR acquisition-based recording

**DOI:** 10.64898/2026.11.17.767295

**Authors:** Andrea Fernández-Gómez, Pablo Guridi-Fernández, Dolores L. Guzmán-Herrador, Ivana González-Fernández, Nathan Barrabes, Raphael Laurenceau, Uli Klümper, David Bikard, Matxalen Llosa

## Abstract

Horizontal gene transfer mediated by conjugative plasmids is a primary driver of bacterial evolution and the spread of adaptive traits, including antibiotic resistance determinants. Tracking how these plasmids disseminate through complex microbial communities remains a major challenge. Current methodologies often yield static, endpoint snapshots of host identity, overlooking transient transfer events and plasmid encounters with the co-resident mobilome. To overcome these limitations, we introduce SPacer Integration (SPI) plasmids, which turn the mobile element itself into a recorder of its own transfer history. We engineered conjugative and mobilizable plasmids to carry a CRISPR-Cas adaptation cassette (*cas1–cas2* and a minimal CRISPR array) and maintain tight repression of *cas1-cas2* expression in the donor cell, so that upon transfer they capture spacers from the chromosome and plasmids of new hosts and store them permanently in their own array. Subsequent PCR amplification and sequencing of these expanded arrays provide a molecular record of both the bacterial hosts and the co-resident plasmids encountered. We created SPI plasmids using the backbones of well-characterized mobilizable and conjugative plasmids RSF1010, RP4, F, and R100-1. We tested SPI plasmids in defined laboratory matings and in complex wastewater communities. The analysis of the acquired spacers successfully mapped permissive host ranges and profiled the co-resident plasmids *in situ*. Requiring only standard cloning, PCR and sequencing, this accessible and cost-effective strategy establishes a new framework for studying plasmid ecology. Because spacers are integrated chronologically, it further offers the potential to reconstruct multi-step transfer trajectories across natural microbial networks.

## Introduction

Horizontal gene transfer (HGT) is a major driver of bacterial adaptability, enabling the rapid acquisition of beneficial traits across lineages (Arnold et al., 2021). This capacity for genetic exchange is particularly significant within natural microbial communities, where dense and taxonomically diverse populations create opportunities for gene flow between unrelated species (Brito, 2021). Among the mechanisms mediating HGT, conjugation, the direct cell-to-cell transfer of plasmid DNA mediated by Type IV Secretion Systems, is especially efficient at disseminating adaptive traits, including antimicrobial resistance genes, across both clinical and environmental settings (San Millan, 2018). Broad-host-range (BHR) plasmids are of particular concern in this context, as they can cross taxonomic boundaries and introduce resistance determinants into phylogenetically distant hosts (Coluzzi and Rocha, 2025). Understanding how conjugative and mobilizable plasmids move through natural communities is therefore essential both for elucidating the mechanisms underlying HGT and for informing strategies to limit the spread of clinically relevant genes.

This has motivated the development of a range of strategies to track plasmid dissemination in complex communities. Early approaches relied on tagging plasmids of interest with selectable or visible markers, such as antibiotic resistance genes or fluorescent proteins, allowing the identification of transconjugants and enabling measurements of plasmid transfer in complex environments (reviewed in Sørensen et al., 2005; Brito, 2021). However, reporter-based approaches can be skewed by resistances already present in the community, or typically depend on cultivation, thus excluding the vast uncultured majority. More recently, an approach that tags plasmids with a catalytic RNA to permanently barcode the 16S rRNA of recipient cells upon transfer has enabled culture- and translation-independent transconjugant identification, creating a direct and persisting molecular link between plasmid uptake and host taxonomy (Kalvapalle et al., 2025). Other culture-independent strategies have instead relied on sequencing and computational analysis: bulk metagenomics can catalog the plasmids present in a community but struggles to link them to specific hosts due to assembly fragmentation, while techniques such as proximity ligation (Hi-C), methylation profiling, and single-cell approaches have enabled the association of plasmids with their bacterial hosts (Saak et al., 2020; Brito, 2021). Despite these advances, most current approaches are often experimentally elaborate and are limited to static, endpoint host identification. Consequently, they fail to capture the true dynamics of plasmid dissemination, overlooking transient events that might happen in the community. Capturing this kind of information requires methods capable of recording transfer events as they occur.

One process naturally suited to this kind of real-time recording is CRISPR adaptation, which has recently been repurposed as a tool for programmable molecular recording (Sheth and Wang, 2018). This application exploits the natural ability of CRISPR-Cas systems to sample DNA fragments from the intracellular environment and stably integrate them as spacers into the CRISPR array, a process catalyzed by the highly conserved Cas1–Cas2 complex (Lee and Sashital, 2022). Some properties of this natural process make it particularly well suited for recording applications. First, because new spacers are integrated unidirectionally at the leader-proximal end of the array, allowing to reconstruct past events chronologically. Second, once integrated, spacers are stably and permanently inherited, providing a durable record rather than a transient signal. Third, acquisition occurs independently in each cell, so that a population of recording cells can capture the heterogeneity of the underlying process at single-cell resolution rather than reporting only a population average. This led to the proposal that, if spacer acquisition could be induced, the resulting spacer content could serve as a high-resolution readout of the intracellular environment, one that could be linked to defined environmental or biological triggers (Shipman et al., 2016; Sheth et al., 2017).

The Type I-E CRISPR-Cas system of *Escherichia coli*, the best-characterized Class 1 system, has become the main platform for translating this concept into working tools (Sheth and Wang, 2018). A key feature for recording applications is that spacer acquisition by this system is naturally repressed under standard laboratory conditions. However, overexpression of *cas1* and *cas2* from an inducible plasmid is sufficient to activate acquisition into a CRISPR array present in the cell (Lee and Sashital, 2022). Previous work has shown that the full CRISPR array is not required for spacer acquisition. Instead, a minimal array comprising only the 60 nucleotides of the leader upstream of the first repeat, together with a single repeat, is sufficient. Transcription of the array is not necessary either (Yosef et al., 2012). Moreover, Type I arrays have been shown to be functional when cloned into plasmids (Díez-Villaseñor et al., 2013; Goren et al., 2016; Shipman et al., 2016). The *E. coli* Type I-E adaptation machinery also relies on broadly conserved host factors, such as the Integration Host Factor (IHF), for ordered spacer integration (Nuñez et al., 2016; Kamashev et al., 2017).

This inducible acquisition strategy based on the *E. coli* Type I-E machinery has been applied to record diverse biological and synthetic signals (Shipman et al., 2016, 2017; Sheth et al., 2017; Schmidt et al., 2018, 2022; Yim et al., 2021), including the direct monitoring of HGT within complex, natural bacterial communities (Munck et al., 2020). An engineered *E. coli* recording strain overexpressing Cas1–Cas2 was shown to acquire spacers from plasmids, phages, and other mobile genetic elements (MGEs) upon their entry into the cell, thereby generating a record of HGT events occurring in situ, without relying on selectable phenotypes. However, because this recording capacity resides in a fixed *E. coli* strain, this approach is limited to reporting on DNA that reaches that single recipient, missing transfer events between other community members. It also fails to capture the specific MGEs a plasmid co-resides with, an essential piece of information since such co-occurrences shape its transfer dynamics and evolutionary fate (Brockhurst et al., 2019; Garcillán-Barcia et al., 2025).

Here, we generate SPI (SPacer Integration) plasmids, which are conjugative or mobilizable plasmids of interest engineered to carry their own CRISPR adaptation module, enabling them to acquire spacers in real time from the chromosome and plasmids of their hosts throughout their movement across a community (**Fig. 1a**). By acting as self-recording elements, SPI plasmids allow us to uncover their complete host range, capturing both stable and transient transfer events, while simultaneously revealing the plasmids that they co-reside with. The performance of this system was first characterized using defined donor and recipient strains, and the system was then deployed in natural bacterial communities, revealing previously inaccessible insights into the dynamics of HGT.

**Figure 1.**
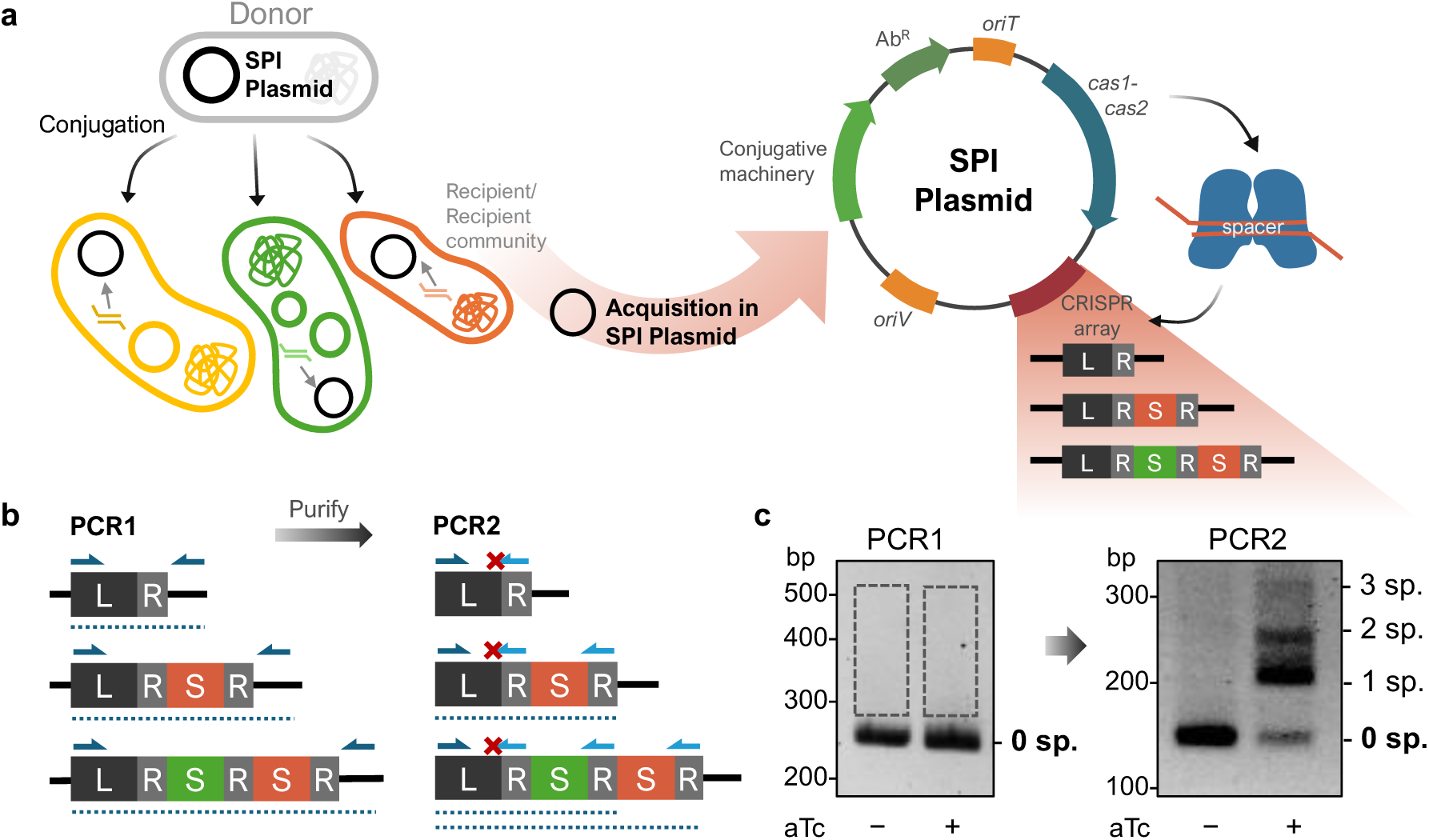
SPI plasmid recording strategy and CAPTURE PCR enrichment. **a** Schematic representation of the SPI plasmid concept. A donor cell transfers the SPI plasmid to diverse recipients within a community by conjugation. Once inside a permissive host, the plasmid-encoded Cas1–Cas2 complex acquires new spacers from the intracellular environment of the host (chromosome or plasmids) and integrates them into the SPI’s minimal CRISPR array. **b** Principle of the CAPTURE PCR strategy used to selectively amplify expanded arrays. After an initial PCR spanning the entire array region (PCR1) and gel purification of the expected size range, a second, enrichment PCR (PCR2) uses a repeat-binding reverse primer whose 5′ end is mismatched to the leader sequence, preventing amplification of non-expanded arrays. **c** Application of CAPTURE PCR to SPI RSFK. Gels show the PCR1 (left) and PCR2 (right) products from a serial-passage assay of *E. coli* carrying both SPI RSFK and pTetR, with (+) or without (−) induction of *cas1–cas2* expression. Rectangles in PCR1 mark the size-selected regions that were excised and purified to serve as templates for PCR2. Molecular weight markers (bp) are shown on the left, and the sizes corresponding to non-expanded (0 sp.) and expanded arrays carrying 1, 2 or 3 spacers are indicated on the right.

## Results

### Generation and validation of SPI RSFK

Our recording strategy builds on previous approaches in which overexpression of the *E. coli* Type I-E Cas1–Cas2 complex has been used to drive spacer acquisition for molecular recording purposes (Munck et al., 2020). We selected the mobilizable plasmid RSFK, derived from RSF1010, as the backbone for our SPI plasmid due to its BHR, high promiscuity, and ability to be mobilized by numerous plasmid groups, including IncP and IncF plasmids (Loftie-Eaton and Rawlings, 2012). To engineer RSFK into a SPI plasmid, we introduced an adaptation module comprising the *E. coli cas1–cas2* genes under the aTc-inducible P*tet* promoter, the *tetR* repressor, and a minimal CRISPR array consisting of the terminal 60 bp of the leader sequence and a single repeat, both derived from the *E. coli* K-12 CRISPR array I (Yosef et al., 2012) (**Fig. 1a**).

We tested whether the resulting plasmid, SPI RSFK-TetR, could acquire spacers into its own array in *E. coli*. The plasmid was introduced into *E. coli* BL21, a *cas*-deficient strain in which acquired spacers cannot trigger interference, and cultures were serially passaged for three rounds with or without induction. Array expansion was assessed using the CAPTURE PCR strategy (McKenzie et al., 2019) (**Fig. 1b**). An initial PCR was used to amplify the array region, followed by excision of the gel region corresponding to the expected size of expanded arrays and purification of the recovered DNA. A second, enrichment PCR is then performed using a repeat-binding reverse primer mix whose 5′ end is deliberately mismatched to the native leader–repeat junction sequence, thereby selectively amplifying expanded arrays at the cost of excluding any spacer that coincidentally ends with the same nucleotide as the leader. PCRs were performed as described using the culture pellet from the third passage, and visualized in an agarose gel (**Fig. 1c**). Induced cultures showed bands corresponding to arrays expanded by one, two, or three spacers, which were absent in the non-induced control, confirming that SPI RSFK-TetR acquires spacers upon induction of the *cas1-cas2* operon.

We next assessed whether the system could record spacers from the host following conjugation. To ensure that spacers were acquired only after transfer into a recipient and not in the donor, where acquisition would be uninformative and could saturate the array, we implemented a zygotic induction strategy: *tetR* was removed from the SPI plasmid (generating SPI RSFK) and instead supplied constitutively from a separate pTetR plasmid stably maintained in the donor. Under this design, TetR represses the adaptation module only in the donor, but upon conjugative transfer into a recipient the module is derepressed and adaptation is activated. As donor strain we used MFDpir, a diaminopimelic acid (DAP)-auxotrophic strain, which allows an easy counterselection of the donor after conjugation by not adding DAP. It also carries an integrated RP4 plasmid, capable of mobilizing RSFK.

Using this setup, we performed conjugations with *Klebsiella oxytoca* or *Klebsiella pneumoniae* as recipients on solid LB medium. After mating, the conjugation mixture was grown on selective LB broth permitting growth of transconjugants only, and culture pellets were used as template for CAPTURE PCR. Expanded arrays were purified by gel extraction and individual arrays were sequenced using Nanopore technology. The obtained reads were processed with custom software to extract spacers, which were then assigned by alignment to their most likely origin (donor, recipient, or plasmids), all of which had available sequence data. The average distribution of spacer origins is shown in **Fig. 2a**, while the individual charts for each independent experiment are provided in **Suppl. Fig. 1**. For both recipients, an average of 70% of spacers originated from the recipient genome, confirming that SPI RSFK acquires spacers efficiently upon entry into a new host. A minor fraction mapped to the donor, likely reflecting residual leaky expression of *cas1* and *cas2* from the P*tet* promoter, while around 15% mapped back to the SPI plasmid itself, consistent with plasmids being preferential substrates for spacer acquisition (Levy et al., 2015) and with the SPI being able to self-acquire once transferred to the recipient. An evaluation of potential cross- mapping between donor and *Klebsiella* genomes revealed that ambiguous spacers accounted for only 3.28 ± 1.45% and 5.32 ± 1.36% of the *K. oxytoca* and *K. pneumoniae* datasets, respectively, confirming the reliability of our recipient-specific assignments. These results confirm that the system reliably records transfer events and that zygotic induction strongly favors acquisition in the recipient.

**Figure 2.**
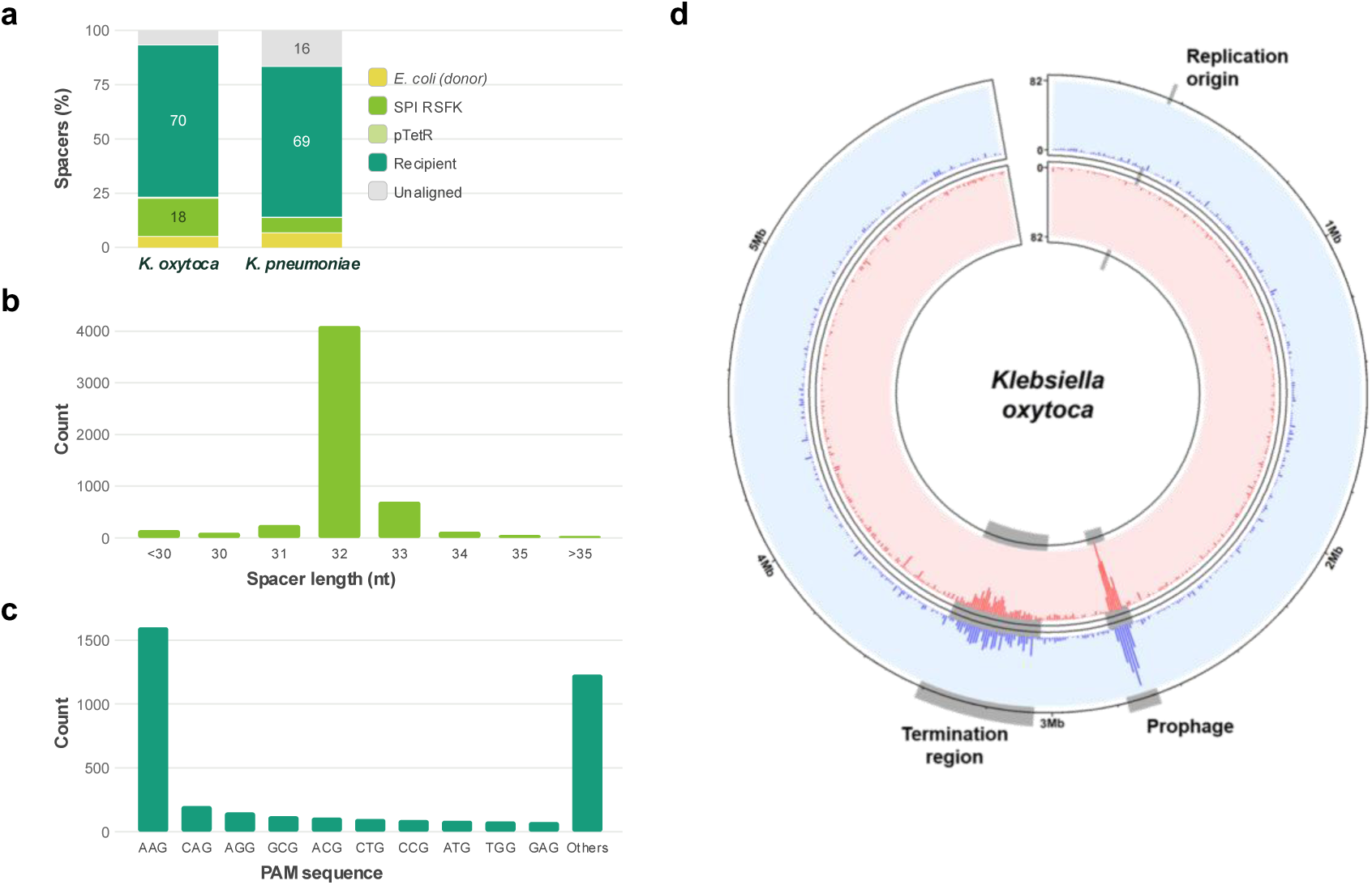
Characterization of spacer acquisition during conjugations of SPI RSFK to *K. oxytoca* and *K. pneumoniae*. **a** Origin of acquired spacers during conjugation from the *E. coli* MFDpir donor to *K. oxytoca* and *K. pneumoniae*. The bar chart show the mean percentage of spacers mapping to each genome or plasmid origin (averaged from three independent experiments for *K. oxytoca* and two for *K. pneumoniae*). **b, c** Combined distribution of spacer lengths (b) and PAM consensus sequences (c) pooling all replicates for conjugations to both recipients. **d** Chromosomal mapping of the spacers acquired in conjugations to *K. oxytoca* (combined from three replicates) using a 10,000 bp window size. The outer blue ring and inner red ring represent acquisition from the positive and negative strands, respectively. Peak heights denote the number of acquired spacers, with the vertical axis at position 0. Key regions are highlighted.

We verified that the adaptation module of the plasmid retained the canonical features of the native Type I-E adaptation machinery. The mean spacer length was 32 bp, as expected (**Fig. 2b**), and the preferred PAM sequence was AAG (**Fig. 2c**), although present in fewer than half of spacers, consistent with the low PAM specificity reported for this system (Mitić et al., 2023). Finally, we sought to determine from which regions of the recipient genome spacers were being acquired, using *K. oxytoca* as a model. Mapping the spacers derived from its chromosome back onto the *K. oxytoca* genome (**Fig. 2d**) revealed broad, near-uniform coverage across the chromosome and between the two DNA strands, with two exceptions corresponding to the replication terminus region and a prophage, both previously described as natural spacer acquisition hotspots (Levy et al., 2015; Nobrega et al., 2020). Together, these results demonstrate that SPI RSFK is fully functional, capable of acquiring spacers following conjugative transfer, and preserves the preferences of the Type I-E adaptation system.

## Tracking SPI RSFK host range in a wastewater community

Having characterized the recording system using a single donor, we next asked whether SPI RSFK could record its own transfer history within a natural, complex bacterial community. For that we chose as recipient a wastewater community obtained from the influent of a water treatment plant in Dresden, Germany, enriched by growth in LB to increase overall bacterial density and favor enterobacteria, a group well represented in reference databases and therefore well suited for a proof-of-concept assessment.

Conjugation to the wastewater community was performed by mixing donor and recipient community at a 1:1 ratio based on OD₆₀₀, using a filter-mating setup to ensure efficient cell-to-cell contact, and allowing mating to proceed for 24 h. Total DNA was extracted directly from the recovered conjugation mixture and used as template to amplify the arrays via CAPTURE PCR.

We analyzed the sequenced arrays to extract spacers, which were first filtered against the SPI RSFK and pTetR sequences to remove their associated spacers (**Suppl. Fig. 2a**). Because the identity of the recipients was not known, the remaining spacers (82%) were assigned taxonomically by BLAST combined with a lowest common ancestor (LCA) algorithm, using a reference database of bacterial genomes and plasmids. Since donor-derived spacers cannot be reliably distinguished from those acquired from *E. coli* already present in the community, donor and recipient *E. coli* were considered jointly throughout this analysis. Results are shown in **Fig. 3**. Spacers were assigned across 65 strains spanning 27 genera, all within the phylum Pseudomonadota, consistent with the known host range of the backbone RSF1010 plasmid (Meyer, 2009). Acquired spacers predominantly derived from three Gammaproteobacteria families, Aeromonadaceae, Enterobacteriaceae, and Pseudomonadaceae, with *E. coli*, *Klebsiella pneumoniae* and *Pseudomonas aeruginosa* as the most abundant species, alongside substantial contributions from several *Aeromonas* species. A small fraction of spacers mapped to Betaproteobacteria. This distribution matches the expected composition of an LB-enriched wastewater community and demonstrates that the Cas1–Cas2 adaptation machinery remains functional across phylogenetically diverse recipients spanning multiple bacterial orders, supporting the robustness of SPI plasmids as a recording tool in heterogeneous communities.

**Figure 3.**
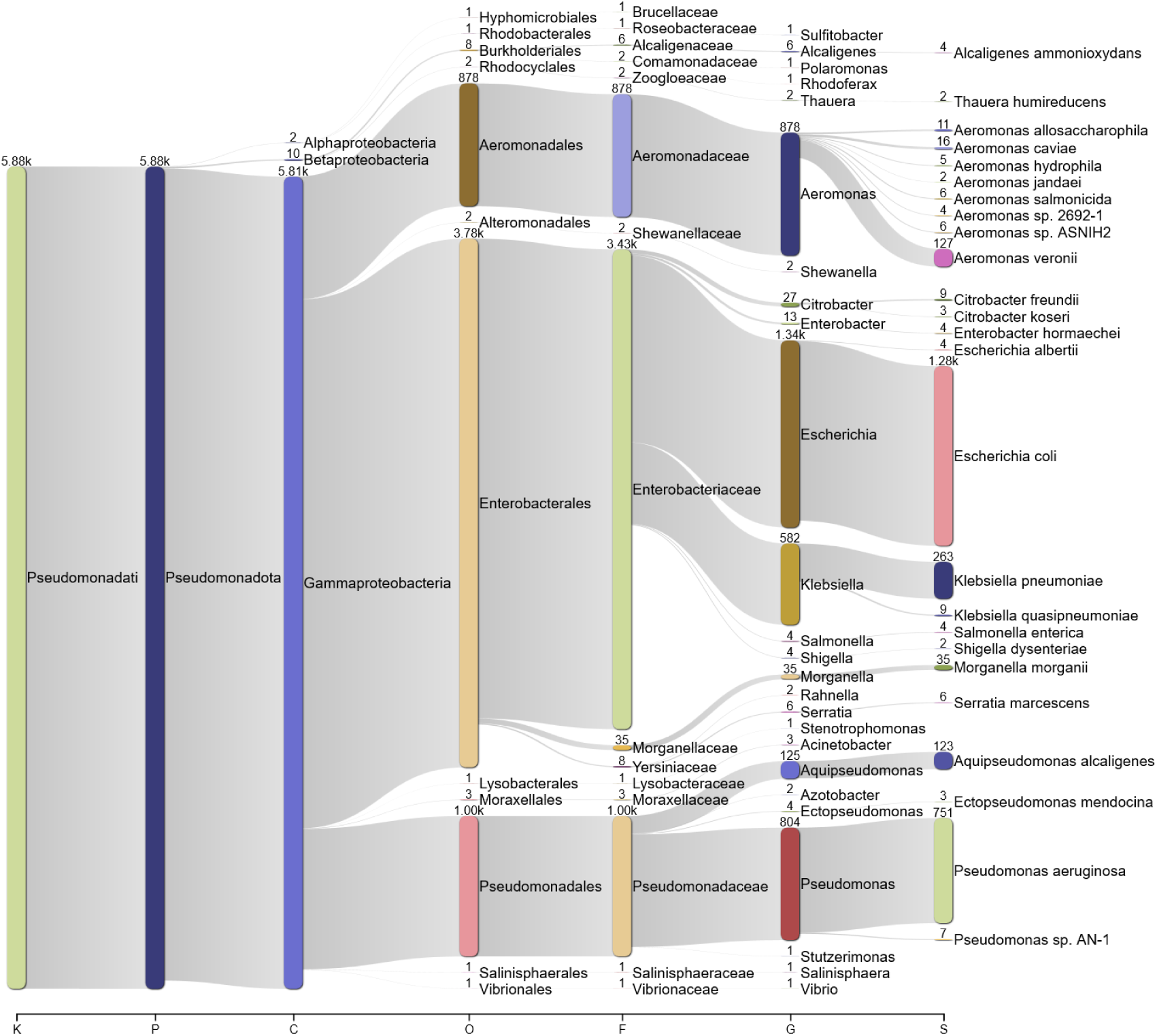
Taxonomic profiling of spacers acquired by SPI RSFK in a complex wastewater community. The diagram illustrates the taxonomic classification of the community-derived spacers from the conjugation of SPI RSFK to the wastewater sample, obtained from four independent sequencing runs. It spans taxonomic ranks from kingdom (K) down to species (S), displaying the 25 most abundant taxa per rank. Taxonomic assignments were performed using an LCA algorithm applied to BLAST hits sharing the highest bitscore.

Because each conjugation is dependent on the specific microbial community present, we further tested SPI RSFK by conjugating it into a new wastewater sample obtained from the influent of a treatment plant in Lloredo, Spain. To assess the impact of culture bias and observe potential differences with an unenriched sample, matings were performed in parallel using both the LB- enriched sample and the raw wastewater sample, which was concentrated to increase bacterial density and promote cell-to-cell contact. The taxonomic profiles of the acquired spacers for the enriched and unenriched conditions are shown in **Suppl. Figs. 2a** and **2b**, respectively. While the main bacterial families were maintained across both settings, the resulting profiles exhibited differences: genera such as *Alcaligenes* and *Aeromonas* were more represented among the spacers from unenriched wastewater, as they can be outcompeted by fast-growing enterobacteria during LB enrichment. To test if spacer acquisition was strictly attributable to conjugative transfer and to assess the potential contribution of other HGT mechanisms, a conjugation-deficient negative control (SPI RSFK oriT-) was tested in parallel with the LB-enriched wastewater mating. As expected for a non-mobilizable SPI plasmid whose adaptation module is repressed in the donor, spacer acquisition was highly reduced, yielding only six spacers in total: three returning no valid BLAST hits, two mapping to the SPI plasmid, and one to *Aeromonas*. While this single environmental spacer indicates that other HGT mechanisms might occasionally occur at practically negligible frequencies, the drastic reduction in spacer uptake by the *oriT*- control demonstrates that the system mainly records conjugative events. Altogether, these results confirm that the system acquires spacers from the available pool of recipients in a conjugation-dependent manner.

To confirm that the taxa identified through spacer acquisition were genuinely present in the community, and to explore whether acquisition frequency reflected host abundance, we sequenced the 16S rRNA gene from the same DNA used for array amplification. Because 16S sequencing resolves taxonomy to the species level, spacers were reassigned for this specific comparison using their best BLAST hit, without LCA filtering, to allow direct species-level comparison with the 16S data. The relationship between the two datasets is shown in **Fig. 4** for the German wastewater sample, and in **Suppl. Fig. 3** for the Spanish ones. Nearly all taxa identified from acquired spacers were independently detected by 16S sequencing, confirming that spacers originated from community members present in the sample. However, the opposite was not true: several abundant community members were absent from the spacer pool, most notably Gram-positive taxa such as *Lysinibacillus* in the case of the sample from Germany, which dominated the 16S profile but did not appear among recipients, consistent with their expected inefficiency as conjugation recipients for this plasmid. Conversely, some less abundant Gram-negative taxa, such as *K. pneumoniae* and *P. aeruginosa*, were largely represented among acquired spacers relative to their 16S abundance, reflecting their efficiency as recipients. Together, these results show that spacer acquisition frequencies reflect the intrinsic permissiveness of each taxon to the SPI plasmid and confirm that the SPI system provides a reliable approximation of the host range of a plasmid within a given community, achieved solely through PCR and sequencing.

**Figure 4.**
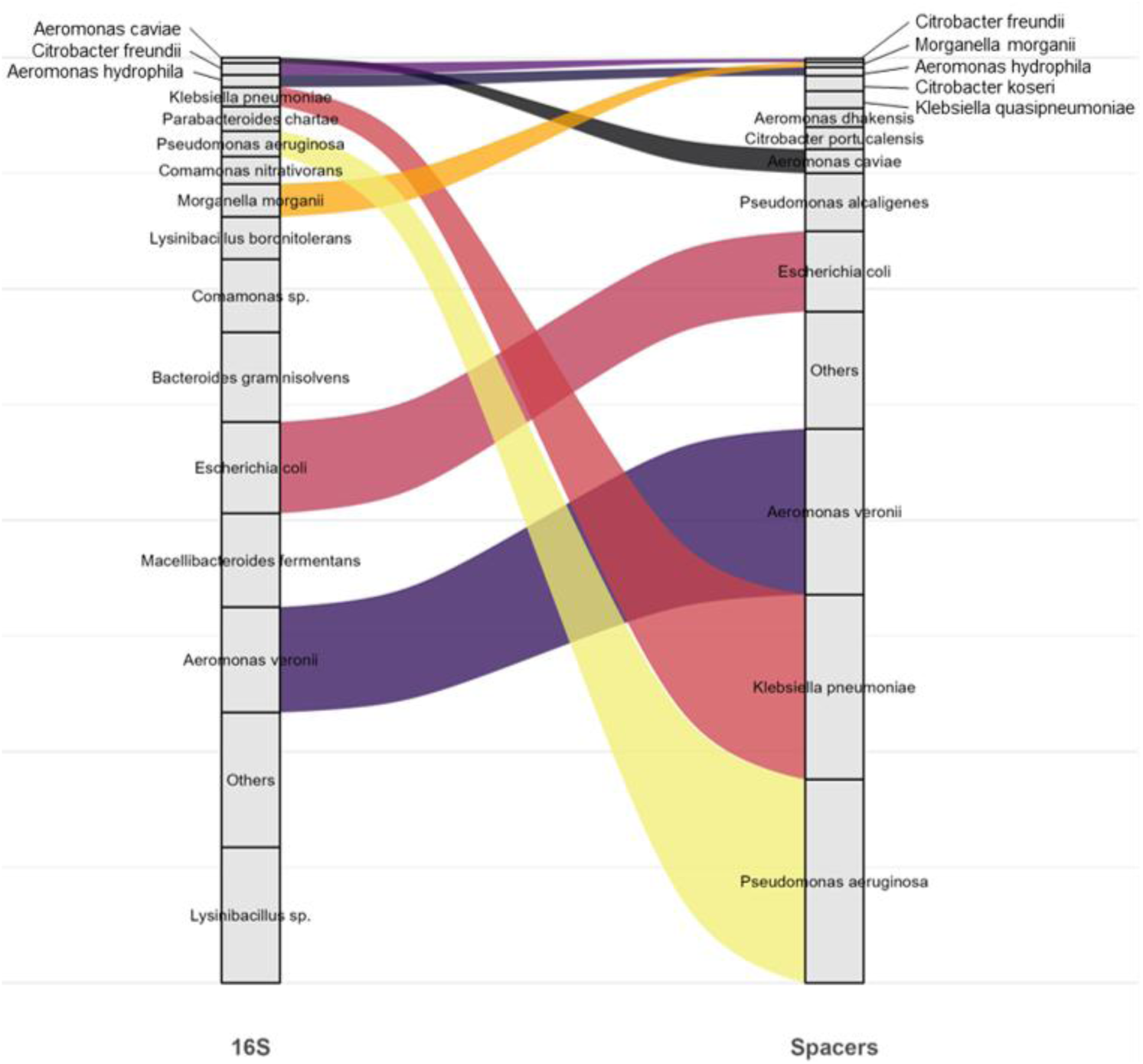
Correlation between microbial community composition and spacer acquisition. Alluvial diagram comparing the relative abundance of bacterial species in the wastewater community, determined by 16S rRNA sequencing (left), with the frequency of acquired spacers assigned to those species via Best BLAST Hit (right). Flow widths are proportional to the relative frequency of each taxon across both datasets. The *Escherichia coli* category includes the donor strain. Species outside the top 20 most abundant are grouped as ‘Others’.

Beyond profiling the overall host range, a unique and valuable application of the SPI system would be tracking the transfer trajectories of individual plasmid molecules across multiple hosts, which could theoretically be reconstructed from arrays acquiring two or more spacers from successive hosts. We attempted to trace these transfer events in SPI RSFK matings. However, arrays carrying multiple spacers proved to be extremely scarce, and PCR modifications aimed at enriching these longer arrays promoted the generation of chimeric PCR amplicons that were unreliable for trajectory reconstruction. Overcoming these limitations to reliably capture multi-host trajectories is a current goal, as it would provide novel and unique information distinctive of the SPI system.

## Co-residence of SPI RSFK with the wastewater mobilome

Natural bacterial communities harbor a large and diverse pool of plasmids, which also constitute a source for CRISPR spacers acquired by adaptation systems (Levy et al., 2015; Androsiuk et al., 2024). Having shown that SPI RSFK records the identity of its recipient cells, we next asked whether it also captures its plasmid content by acquiring spacers directly from these co-resident elements.

To distinguish between chromosomal and plasmid sources, we compared results from two independent BLAST searches: the one used throughout the previous analyses against a database combining bacterial genomes and plasmids, and a second of only plasmid sequences. Spacers with hits only in the combined genomes and plasmids database were classified as chromosomal, while spacers with hits in the plasmid-only database were classified as plasmid-derived if they had no corresponding hit against a chromosome in the first search, or as shared between both genomic contexts if they did. The resulting distribution of spacer origins for the different conjugations to wastewater is shown in **Suppl. Fig. 4a**. Plasmid-exclusive and shared spacers together accounted for 40-60% of all classified spacers, confirming the expected preference of the adaptation machinery for plasmid targets and the substantial degree of sequence sharing between plasmids and chromosomes. To validate this classification, we examined the functional annotation of the genes targeted by spacers in each category. **Suppl. Fig. 4b** shows the result for a representative example. As expected, chromosome-exclusive spacers were enriched in core housekeeping functions such as metabolism and transport, plasmid-exclusive spacers were enriched in accessory and self-maintenance functions such as replication and conjugation, and shared spacers showed a hybrid signature and mobile-element functions, consistent with their dual chromosomal and plasmid distribution.

Since both the plasmid-exclusive and the shared spacers correspond to sequences that can be found on plasmids, we combined these two categories for the analysis of the specific plasmid groups that SPI RSFK had encountered in the three performed matings. The plasmids targeted by the spacers were classified via the COPLA tool (Redondo-Salvo et al., 2021) into plasmid taxonomic units (PTUs), defined as clusters of plasmids sharing high nucleotide identity and evolutionary history that function as natural "molecular species". Because a single spacer sequence could match plasmids belonging to different PTUs, we calculated a weighted presence score for each PTU, defined as the mean fraction of hits directed to that PTU across all spacers with plasmid hits. Plasmids that do not belong to a sufficiently large cluster remain unassigned to any PTU. The resulting PTU profiles are shown in **Fig. 5**.

**Figure 5.**
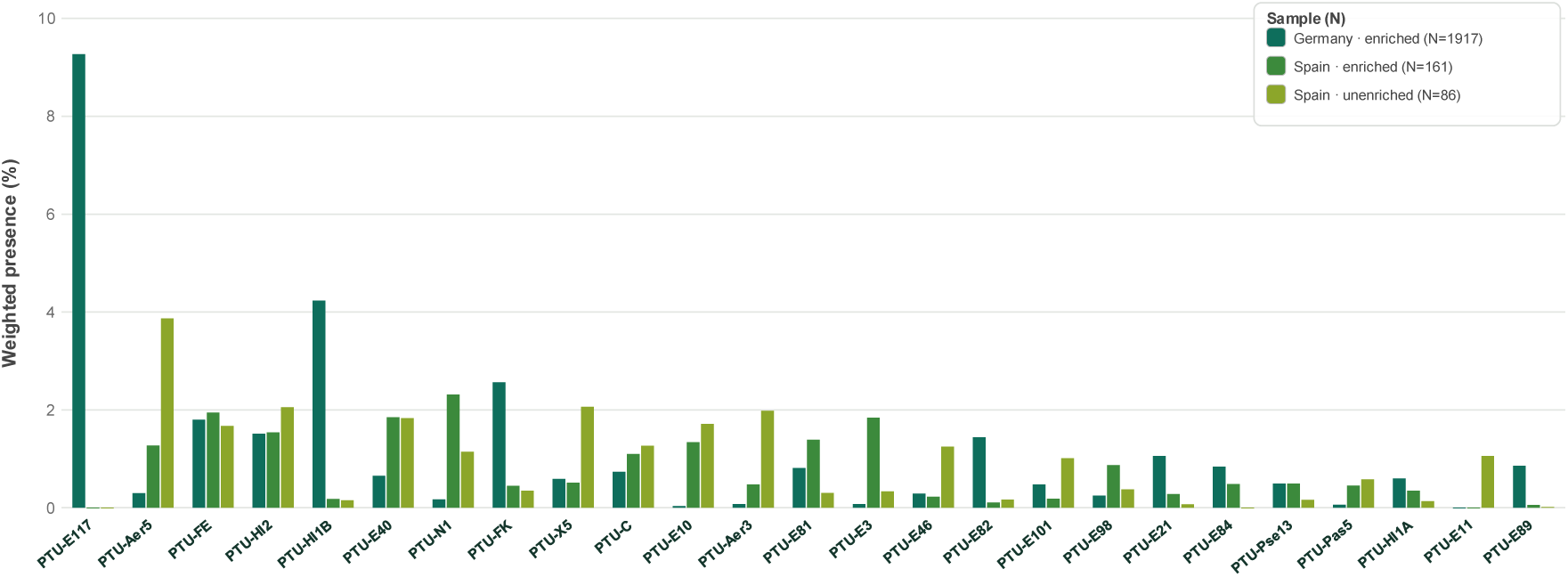
**PTU classification of plasmid-derived spacers acquired by SPI RSFK in wastewater communities**. Weighted presence of the 25 most abundant PTUs among the potentially plasmid-derived spacers, shown for the three wastewater matings as parallel bars. This score is a normalized measure in which each spacer contributes a total of one, distributed across its PTU-assigned BLAST hits and averaged over the number of spacers with at least one PTU-assigned hit (N, indicated in the legend). Candidate PTUs and provisional PTUs were excluded from the plot. The 25 PTUs with the highest summed weighted presence across the three samples are shown. The Germany sample combines four independent sequencing runs, the LB-enriched Spain sample two, and the unenriched Spain sample a single run.

Across the three wastewater samples, the detected PTU profiles were consistent with the known plasmidome of Enterobacterales, the most prevalent order among the transconjugants. Abundant groups included, among others, PTU-FK and PTU-FE, large conjugative narrow-host- range (NHR) plasmids frequently associated with resistance and virulence genes in *Klebsiella* and *Escherichia*, respectively; PTU-HI1B and PTU-HI2, conjugative plasmids distributed across multiple enterobacterial genera, with PTU-HI2 in particular being one of the most antibiotic resistance gene-laden PTUs described; and PTU-C, a BHR group also strongly associated with AMR. Overall, the recovered PTUs spanned a wide range of plasmid lifestyles and functions: alongside these conjugative, often resistance- or virulence-associated groups, we also detected additional conjugative groups (e.g. PTU-N1, PTU-X), mobilizable and non-mobilizable groups carrying no known resistance determinants, and even *Aeromonas*-associated environmental PTUs (e.g. PTU-Aer3, PTU-Aer5) that fall outside the enterobacterial plasmidome. Among the non- mobilizable groups was PTU-E117, the most abundant PTU in the German wastewater dataset. Notably, some of the detected PTUs, including PTU-FE, PTU-FK or PTU-X, are known to mobilize RSF1010-type plasmids. Their presence among the recipients suggests that SPI RSFK could have been secondarily mobilized and disseminated further through the community, beyond the initial donor-to-recipient transfer, highlighting the importance of identifying potential multi-step transfer trajectories. Together, these results demonstrate that the SPI plasmid system can sample, *in situ*, the portion of the community mobilome with which it co-resides, spanning both well-characterized and relevant plasmid groups and other, less-studied ones that remain to be further explored.

## Extending the SPI platform to conjugative plasmids

We extended our approach to generate conjugative SPI plasmids which do not rely on a compatible helper for mobilization, allowing them to disseminate autonomously across microbial communities and potentially capture more transfer events. We selected three well-characterized conjugative plasmids: RP4, a PTU-P1 plasmid widely used as a model for its BHR (Le et al., 2025); pOX38, a derivative of the *E. coli* F plasmid (Guyer et al., 1981); and R100-1, a NHR PTU-FE plasmid that is one of the earliest described multidrug-resistance elements (Egawa and Hirota, 1962). Each of these plasmids was converted into a SPI by integrating into their backbones the adaptation module, *cas1–cas2* under the P*tet* promoter, together with the minimal CRISPR array, omitting the *tetR* repressor. These conjugative SPIs were therefore maintained and propagated in host strains carrying a separate pTetR plasmid supplying TetR in trans, analogous to the strategy used for SPI RSFK.

To evaluate the performance of these conjugative SPI plasmids, we tested all three in matings with the wastewater community from Spain, in both the LB-enriched and unenriched conditions. Following the same protocol established for SPI RSFK, total DNA was extracted directly from the conjugation mixtures and subjected to CAPTURE PCR to amplify expanded arrays. The resulting amplicons were sequenced, and the origin of the newly acquired spacers was determined. The proportion of recovered spacers mapping to the SPI plasmid, the pTetR helper, or remaining unassigned and thus taken forward as potential recipient-derived is shown in **Suppl. Fig. 5a.** The total number of recovered spacers showed a high variability between plasmids and conditions, ranging from < 200 to >2000, preventing us to use it as an estimation for how efficiently each SPI plasmid acquires spacers. The level of self-acquisition (presence of spacers matching the plasmid itself) also varied among experiments/samples, but RP4 showed a low rate (<25%) while R100-1 and especially pOX38 acquired about 50% of its spacers from its own backbone (up to 82% for pOX38 in the unenriched condition).

The taxonomic profiling of the potentially community-derived spacers was performed as for SPI RSFK, using BLAST combined with an LCA algorithm. Results, shown in **Suppl. Fig. 5b**, revealed that, in all conditions, acquisitions were dominated by a small number of Gammaproteobacterial families: Enterobacteriaceae, Aeromonadaceae and, to a lesser extent, Alcaligenaceae. *Escherichia* was consistently the most abundant genus, reflecting not only its efficient conjugation with all three plasmids but also the fact that this taxon includes the spacers acquired in the donor. The unenriched matings again showed a higher relative contribution of Aeromonadaceae and Alcaligenaceae, taxa that are outcompeted during LB enrichment. Low spacer numbers and limited taxonomic resolution preclude firm conclusions. Nevertheless, the observed profiles align with expectations: the BHR plasmid RP4 (PTU-P1) showed a wider recipient spectrum than pOX38 and R100-1 (PTU-FE), which are mainly confined to Enterobacteriaceae.

Finally, we also investigated the acquisition of plasmid-derived spacers by the conjugative SPI plasmids using the previously described method. The relative proportions of spacers originating from host chromosomes, plasmids, or that could come from either origin for each SPI plasmid and condition are shown in **Suppl. Fig. 6a**. For spacers that could be derived from plasmids, the PTUs of the potential plasmid targets were assigned, and the resulting PTU distributions are presented in **Suppl. Fig. 6b**. Although the overall low numbers of spacers with hits to characterized PTUs limits our ability to establish generalized trends, certain plasmid groups stood out. For instance, spacers targeting PTU-Q1 plasmids were notably prominent in the unenriched matings for both RP4 and R100-1, while hits mapping to PTU-FE were consistently detected across all three SPI plasmids.

Altogether, conjugative SPI plasmids are functional tools that acquire community-derived spacers from both host chromosomes and resident plasmids, recording their host range and intracellular interactions. Multiplexing future experiments to increase spacer yields will provide a deeper understanding of their transfer dynamics.

## Discussion

Reconstructing how plasmids move through natural bacterial communities has traditionally relied on static, endpoint-based methods (Brito, 2021), which reveal where a plasmid has ended up but not the trajectory or intermediate hosts it passed through, nor the MGEs it encountered along the way. Building on the demonstration that CRISPR spacer acquisition can record HGT events entering a recipient (Munck et al., 2020), we inverted this design by placing the recording machinery directly onto the mobile element itself. The SPI Plasmid strategy validated here shows that this shift is both feasible and informative, providing a chronological, self-contained record of the transfer history of a plasmid.

SPI RSFK, the plasmid on which this strategy was primarily developed, proved functional for spacer acquisition in the recipient, first in conjugations to isolated recipient strains and subsequently in complex communities. The zygotic induction strategy, achieved by decoupling TetR from the SPI, was effective at minimizing donor-derived noise in the initial single-recipient conjugations. This consistently low background suggested that donor-derived acquisition would remain similarly low during conjugations to complex communities, and most of the *E. coli* signal detected in the community experiments could be attributed to transfer into native *E. coli* residents, rather than acquisition from the donor. Nonetheless, further reducing this residual donor noise remains a desirable improvement for cleaner future datasets. The RSFK adaptation module also preserved the intrinsic features of its parental Type I-E system, including uniform spacer acquisition across target sequences and the expected PAM preference.

Extending this system to an LB-enriched wastewater community demonstrated, to our knowledge, the first plasmid-based molecular recorder used in an undefined, complex microbial environment. The resulting host-range profile, dominated by Enterobacteriaceae, Pseudomonadaceae, and Aeromonadaceae, closely matches the established host range of RSF1010 among Pseudomonadota (Meyer, 2009) and aligns with independent observations of RSF1010 dissemination in other engineered water systems, such as groundwater-fed rapid sand filters (Pinilla-Redondo et al., 2021), reinforcing that the taxonomic signal captured by the SPI plasmid reflects biological permissiveness and is not an artifact of the recording system. To ensure the robustness of this signal, we applied the system to wastewater samples from different treatment plants, testing both LB-enriched cultures and raw, unenriched water. The SPI system captured the expected compositional shifts driven by sample origin and LB enrichment, recording the permissive hosts available in each specific context. The obtained profiles emerged from unbiased spacer acquisition, and its correlation with independent 16S rRNA profiling confirmed that acquisition frequency does not scale linearly with the abundance of a taxon. Ultimately, the recorded profiles reflect two successive filters: first, the ecological filter of which taxa survive or dominate under the specific conditions (natural or enriched), and second, the mechanistic filter of which bacteria are efficient recipients capable of receiving the plasmid and supporting Cas1–Cas2 expression and spacer acquisition. Notably, spacer acquisition proved functional not only in Enterobacterales, the native order of the chosen *cas1–cas2*, but also in Pseudomonadales, Aeromonadales, and Burkholderiales. To our knowledge, this is the first demonstration that these *E. coli* Type I-E proteins are active outside their native species, and their functionality in Betaproteobacteria is particularly notable given the greater phylogenetic distance involved. This suggests that the host factors required for adaptation, such as IHF (Nuñez et al., 2016), are sufficiently conserved across Pseudomonadota to support this system across the phylum, broadening the applicability of this recording strategy to mixed communities.

Beyond host identity, the high proportion of plasmid-derived spacers recovered from the wastewater datasets is consistent with the well-documented preference of the Cas1–Cas2 complex for plasmid substrates, and the distinct functional signatures separating plasmid- and chromosome- assigned spacers, together with the intermediate, MGE-enriched profile of spacers matching both compartments, further support that the SPI is sampling distinct genomic contexts. Regarding the plasmid-derived fraction specifically, the resulting PTU profile of the putative plasmid targets, dominated by well-characterized, often resistance-associated conjugative groups alongside less- studied elements, is consistent with previously described wastewater plasmidomes (Smyth et al., 2025) and illustrates a capability that, to our knowledge, is not offered by any other current method for tracking HGT in complex communities: the simultaneous capture of host-range information and of the co-resident mobile elements a plasmid has encountered along its trajectory.

To further expand the utility of this approach, we extended our strategy to engineer fully conjugative SPI plasmids representing ecologically and clinically relevant groups. These self- transmissible elements are capable of autonomous dissemination, and we demonstrated their ability to successfully acquire community-derived spacers directly in complex wastewater samples. We observed variability both between independent experiments and among the plasmids themselves, regarding the extent of self-acquisition and the proportion of spacers acquired from resident plasmids versus host chromosomes. Although they also exhibited generally lower overall acquisition rates compared to SPI RSFK, these conjugative SPIs still yield highly valuable information regarding their host range in wastewater samples and interactions with co-resident plasmids. Future applications will aim to scale up or multiplex the conjugation assays to capture larger spacer pools.

In spite of its utility, the SPI approach also faces some limitations. Taxonomic resolution is constrained by spacer length as, at an average length of 32 bp, some spacers could not be confidently assigned or were resolved only to broader taxonomic ranks. Another technical caveat is that because all arrays generated by a given SPI share the same minimal repeat sequence, multi-spacer amplicons are susceptible to template switching during PCR, generating chimeric arrays that would be misread as a single, continuous trajectory. Indeed, this artifact has currently prevented the reliable reconstruction of true multi-host trajectories from our datasets, which can be inferred from multi-spacer arrays with consecutive spacers assigned to different taxa. Nevertheless, we are modifying the experimental protocol to overcome this barrier, as detecting these sequential transfer events would provide an additional, valuable layer of ecological information. Future implementations should consider strategies to minimize these problems, including modifications to the nested PCR approach to prevent template switching, the use of higher-accuracy sequencing platforms to maximize the taxonomic resolution obtainable from the 32-bp spacers, more active Cas1–Cas2 variants that generate broader pools of spacers or alternative inducible promoters, and even multiplexing them to broaden the range of hosts and elements sampled within a single community.

An additional issue that warrants further investigation is the host range over which the SPI plasmid approach is functional. The successful demonstration of spacer acquisition in taxonomically distant proteobacteria represents a significant achievement. However, for SPI plasmids to become a fully functional tool for tracking HGT in complex bacterial communities, their applicability should be tested in more distantly related bacteria, such as Bacteroidetes and Gram- positive species. Experiments are currently underway in our laboratory to assess SPI-mediated spacer acquisition in these bacteria by adapting the SPI backbone with appropriate promoters to drive Cas1-Cas2 expression and IHF production.

Beyond these optimizations, a key strength of the approach is its accessibility: converting a plasmid of interest into a SPI plasmid requires only standard molecular cloning, and the resulting assay relies solely on PCR and sequencing, making the method straightforward to implement and scale. Building on this versatility, future applications can expand beyond wastewater to other relevant microbiomes. Customized SPI plasmids can be engineered from MGEs of ecological or clinical significance and used to track HGT in the complex communities where their dissemination is most impactful, such as soil communities, clinical settings, or the gut microbiome, among others.

Altogether, these results establish SPI Plasmids as a validated strategy for studying plasmid-mediated HGT directly within complex communities. The approach combines capabilities not previously available in a single method: recording horizontal transfer events within the mobile element itself, capturing co-resident mobile elements alongside host identity, and doing so autonomously, without cell sorting, imaging, or prior knowledge of community composition. This system shifts the study of plasmid dynamics from static snapshots toward a deeper, more integrated understanding of how mobile elements disseminate through natural microbial networks.

## Materials and Methods

### Bacterial strains and culture

Bacterial strains used in this work are listed in **Table 1**. All strains were grown at 37 °C in Luria-Bertani (LB) broth, or on LB agar for solid culture, supplemented with the appropriate antibiotics or supplements as required. They were used at the following concentrations: chloramphenicol (Cm), 25 µg/mL; ampicillin (Ap), 100 µg/mL; kanamycin monosulfate (Km), 50 µg/mL; streptomycin (Sm), 300 µg/mL; gentamicin sulfate (Gm), 10 µg/mL; anhydrotetracycline hydrochloride (aTc), 200 ng/mL; and diaminopimelic acid (DAP), 0.3 mM.

**Table 1.**
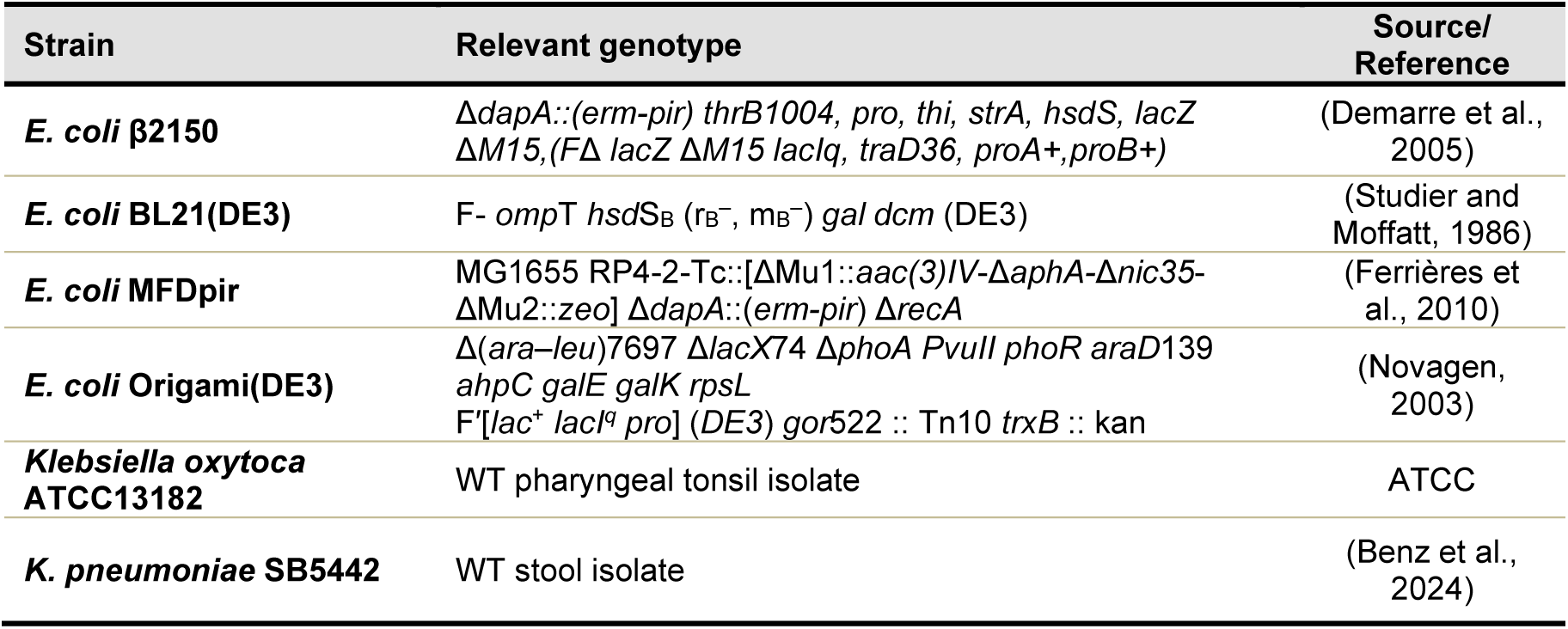
Bacterial strains used in this work.

| **Strain** | **Relevant genotype** | **Source/ Reference** |
|---|---|---|
| ***E. coli* β2150** | *ΔdapA::(erm-pir) thrB1004, pro, thi, strA, hsdS, lacZ ΔM15,(FΔ lacZ ΔM15 lacIq, traD36, proA+,proB+)* | (Demarre et al., 2005) |
| ***E. coli* BL21(DE3)** | F- *omp*T *hsd*S<sub>B</sub> (r<sub>B</sub><sup>−</sup>, m<sub>B</sub><sup>−</sup>) *gal dcm* (DE3) | (Studier and Moffatt, 1986) |
| ***E. coli* MFDpir** | MG1655 RP4-2-Tc::[ΔMu1::*aac(3)IV*-Δ*aphA*-Δ*nic35*-ΔMu2::*zeo*] Δ*dapA*::(*erm-pir*) Δ*recA* | (Ferrières et al., 2010) |
| ***E. coli* Origami(DE3)** | Δ(*ara–leu*)7697 Δ*lacX*74 Δ*phoA PvuII phoR araD*139 *ahpC galE galK rpsL*<br>F′[*lac*<sup>+</sup> *lacI<sup>q</sup> pro*] (*DE3*) *gor*522 :: Tn10 *trxB* :: kan | (Novagen, 2003) |
| ***Klebsiella oxytoca* ATCC13182** | WT pharyngeal tonsil isolate | ATCC |
| ***K. pneumoniae* SB5442** | WT stool isolate | (Benz et al., 2024) |

## Plasmid construction

All plasmids used in this study are summarized in **Table 2**. Novel plasmids constructed for this work were generated using either Gibson Assembly (Gibson et al., 2009) or λ Red-mediated recombination (Datsenko and Wanner, 2000). For both approaches, specific details regarding the construction method and the origins of the vectors, inserts, or integrated fragments are provided in **Suppl. Table 1**, and the primers used for all PCR amplifications are listed in **Suppl. Table 2**. Briefly, for Gibson Assembly, vector backbones and inserts were amplified by PCR prior to the assembly reaction. For λ Red-mediated recombination, the process was carried out by expressing the λ Red recombinase system (Exo, Beta, and Gam) from the pKOBEG plasmid. When constructing SPI plasmids lacking a native *tetR* gene, transformation of the assembly products or the recombination process were performed in a host strain harboring *tetR* in the chromosome (like Origami(DE3)) or a pTetR plasmid.

**Table 2.**
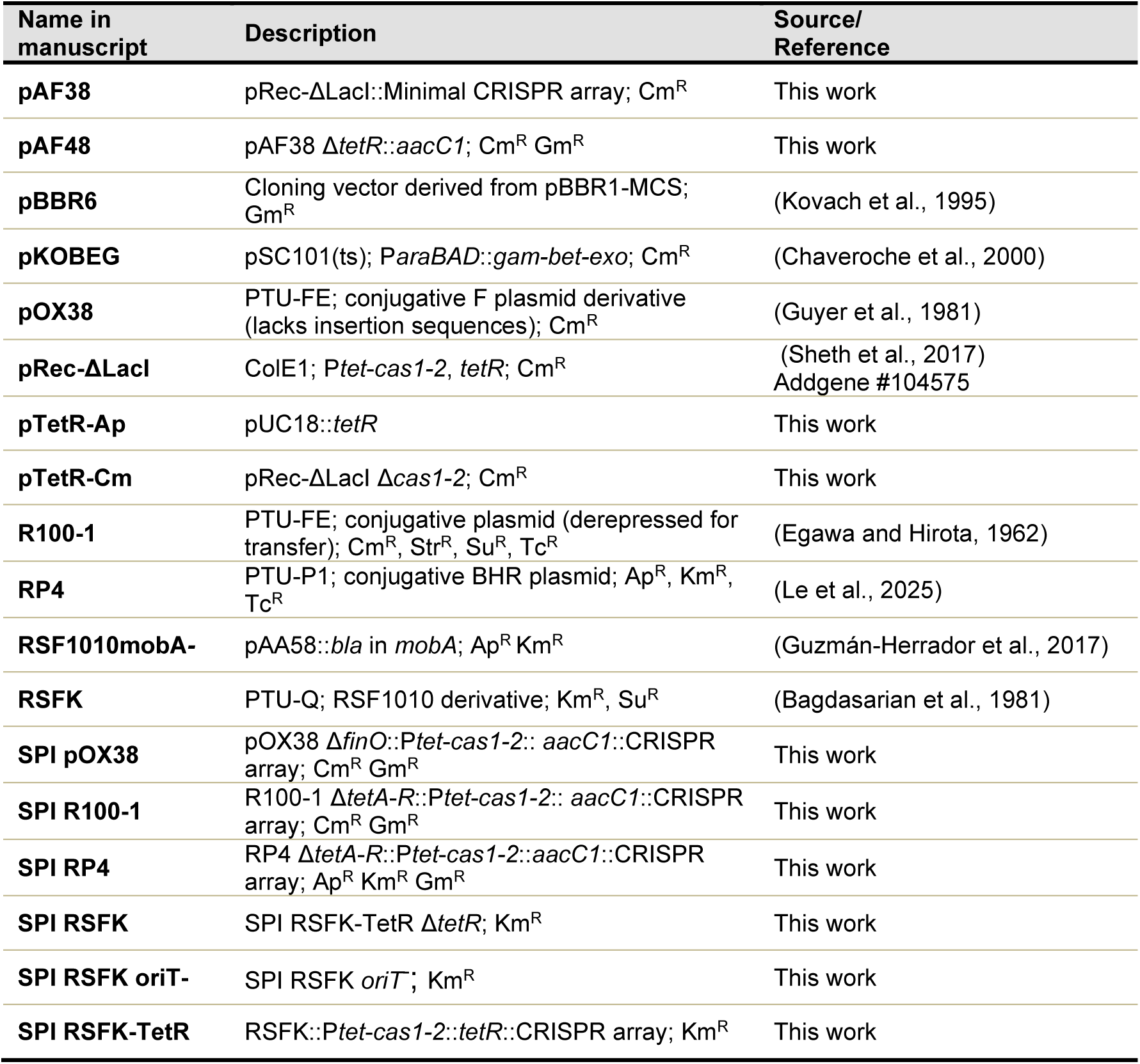
Plasmids used in this work.

## Validation of recording activity by serial passage in *E. coli*

Spacer acquisition by SPI RSFK was first assessed through serial passage of *E. coli* BL21 transformants. Single colonies of the strain carrying the SPI plasmid and pTetR-Cm were inoculated into 5 mL LB medium supplemented with the appropriate antibiotics and aTc to drive *cas1–2* expression, with non-induced cultures maintained in parallel as negative controls. Cultures were grown overnight at 37°C with shaking and diluted 1:300 into fresh medium daily for three successive passages. At each passage, aliquots of the culture were collected, pelleted, and used directly as templates for PCR amplification of the CRISPR array. The resulting PCR products were then run on a 2% agarose gel to detect array expansion.

## Defined conjugation assays

Conjugations of SPI RSFK to defined *Klebsiella* recipients were performed using the DAP- auxotrophic strain MFDpir as the donor, relying on its integrated RP4 system for plasmid mobilization. To control the expression of the system prior to transfer, the donor strain was also transformed with pTetR-Cm plasmid to supply TetR in trans. Donor and recipient strains were grown overnight in LB medium supplemented with the appropriate antibiotics, diluted 1:20 into fresh medium the following morning, and incubated for 3 h. Cells were then harvested by centrifugation, resuspended in fresh LB, and mixed at a 1:1 ratio based on OD, combining 1 OD_600_ unit of each strain. This suspension was concentrated to a final volume of 15 µL and spread evenly across the surface of 24-well plates containing solid LB agar. Mating was allowed to proceed for 2.5 h, after which the mating mixture was recovered in 1 mL of PBS and serially diluted and plated on selective LB agar to enumerate donors, recipients, and transconjugants and calculate conjugation frequencies. In parallel, 10 µL of the recovered mixture were inoculated into 5 mL liquid LB with antibiotics and lacking DAP to select transconjugants, and grown overnight. Cell pellets from this enrichment culture were used for PCR analysis.

## Conjugations to a wastewater microbial community

The complex wastewater microbial communities used as recipients were obtained from the influent of the Kreischa wastewater treatment plant (Dresden, Germany) in September 2025, and from the Lloredo EDAR (wastewater treatment plant, Cantabria, Spain) in July 2026. For the conjugation assays, the preparation of the mating mixtures depended on the recipient condition. For the LB-enriched community, volumes corresponding to 0.5 OD_600_ units of both the donor overnight culture and the enriched recipient community were mixed in 5 mL of PBS. For the unenriched wastewater, 100 µL of the concentrated biomass was mixed with 0.5 OD_600_ units of the donor. The fixed recipient volume was used as an approximation, since the number of viable recipients could not be confirmed before plating (later estimated at 8.4 × 10^6^ CFU/mL). In both cases, the donor and recipient suspension was subsequently passed through a 0.22 µm nitrocellulose filter using a vacuum filtration system to distribute the biomass homogeneously. The filter was then placed onto an LB agar plate and incubated for 24 h. Following the mating period, the mixture was recovered by thoroughly resuspending the cells from the filter in 2 mL of PBS. The recovered mating mixture was processed directly for total DNA extraction. For this step, the MasterPure Complete DNA and RNA Purification Kit was used for the LB-enriched communities, whereas the DNeasy PowerSoil Pro Kit was utilized for the unenriched wastewater samples. In both cases, the resulting purified DNA was used for the subsequent analysis of spacer acquisition.

## 16S rRNA sequencing and community composition analysis

To analyze the composition of the bacterial communities used as conjugation recipients, total DNA extracted from the recovered mating mixture served as a template to amplify the 16S rRNA gene with the primers listed in **Suppl. Table 2**. The resulting amplicons were subjected to Nanopore sequencing via the Microbiome Amplicon Sequencing service provided by Plasmidsaurus, yielding a direct profile of the most abundant species and their relative proportions.

## Nested PCRs for the enrichment of expanded arrays

Array expansion was assessed using CAPTURE PCR (McKenzie et al., 2019), a targeted nested PCR strategy designed to progressively enrich for expanded arrays. PCRs were performed in 20 µL reaction volumes using the NZYTaq II 2x Green Master Mix. For the first-round PCR, the template consisted of either boiled cell lysates from the conjugation mixtures or, in the case of conjugations to the wastewater community, total DNA extracted with the corresponding kit and resuspended in a final volume of 30 µL. This initial amplification used specific primers (**Suppl. Table 2**) to amplify the entire array region of the respective SPI plasmid. To enrich for expanded arrays, a second PCR was performed using 5 µL of the purified first-round product as the template, using the same forward primer paired with a reverse primer engineered to anneal within the CRISPR repeat sequence but with a variable nucleotide at its 3′ end (**Suppl. Table 2**). This mismatch prevents hybridization with the leader sequence, thereby hindering the amplification of unexpanded arrays. Between PCR rounds, size-selective purification was performed to further deplete the unexpanded arrays. This was routinely achieved by electrophoresis on a 2% agarose gel, followed by blind excision of the gel region corresponding to expanded arrays, avoiding the prominent unexpanded array band, and purification with the NZYTech Gelpure kit. Occasionally, size selection with AMPure XP magnetic beads was used instead, initially aiming to retain longer arrays for trajectory detection; however, as long multi-spacer amplicons were found to be prone to PCR chimera formation, only the individual spacer sequences were retained for analysis. The enriched and purified amplicons were submitted to long-read Nanopore sequencing through PlasmidSaurus Premium PCR Amplicon Sequencing service, and the resulting reads were processed using the corresponding software.

## Spacer extraction and assignment pipeline

All custom scripts developed for this work are available in the GitHub repository: https://github.com/fdezgomeza/SPI_plasmids. Raw sequencing FASTQ files were processed using a custom Python (v3.13) pipeline to identify CRISPR repeats in both orientations and extract the intervening spacer sequences. After deduplicating identical arrays, individual spacers were extracted to generate FASTA and FASTQ libraries, preserving array-level linkage and global metrics. For defined donor-recipient experiments, unique spacers were aligned to reference genomes containing the specific donor, recipient, and plasmid sequences using BWA-MEM v0.7.18 (-k 15 -T 15). Potential cross-mapping ambiguities resulting from donor-recipient sequence homology were evaluated using a custom script. For all mapped spacers, the 3-nucleotide sequence immediately upstream of the protospacer target strand was retrieved to evaluate PAM preferences. For experiments involving the complex wastewater community, unique spacers were first aligned against the donor plasmids (SPI plasmid and pTetR) using stringent BWA-MEM parameters (-k 15 -T 24). Unmapped reads, representing potential environmental acquisitions, were extracted and queried against the NCBI RefSeq bacterial database (June 5, 2024 snapshot) and/or a pre-filtered NCBI RefSeq Plasmid database (Release 232, curated to exclude partial, shotgun, and isolated mobile element sequences). Homology searches were performed using BLASTn v2.16.0 (-task blastn-short, E-value ≤ 10⁻⁵, identity ≥ 95%, maximum 50 target sequences). Taxonomic assignment of multi-species hits was resolved using a custom Lowest Common Ancestor algorithm, which assigns taxonomy to the lowest shared rank among hits sharing the highest bitscore. Alternatively, for direct comparisons with 16S rRNA community profiling, a Best BLAST Hit approach (i.e. filtering for the highest bitscore and lowest E-value) was applied to allow species-level assignments.

## Genomic context classification and PTU assignment

To classify the genomic context of the acquired spacers, a BLASTn search was performed against the pre-filtered NCBI RefSeq Plasmid database (Release 232) with 50 hits per query. Spacers previously assigned to the SPI or pTetR plasmids were excluded. Spacer origin was determined by comparing the output of this plasmid-only database with the results from the broader NCBI RefSeq bacterial database (comprising chromosomes and plasmids). Spacers mapping exclusively to the plasmid database were classified as plasmidic, whereas those aligning only with the bacterial database were deemed chromosomal. Spacers returning valid hits in both databases were categorized as ambiguous, belonging to either genomic context. The genes targeted by the spacers were retrieved by identifying overlapping Coding Sequence (CDS) features via the NCBI Entrez API. These genes were subsequently assigned to functional categories using a custom keyword dictionary. For spacers classified as potentially plasmid-targeting, the corresponding target plasmids were assigned to PTU. This was achieved by cross-referencing the BLAST hit accessions with a database, kindly provided by the Garcillán-Barcia laboratory, containing PTU metadata previously determined using the COPLA software (Redondo-Salvo et al., 2021). Target plasmids lacking a designated PTU or absent from this metadata database were excluded. Because a single spacer can map to multiple plasmids belonging to different PTUs, a weighted scoring system was applied to reflect the proportional representation of each PTU. The weighted score for a given PTU was calculated using the following formula:

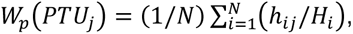

where *N* represents the total number of spacers with at least one hit assigned to a known PTU, *h_ij_* is the number of BLAST hits for spacer *i* assigned to PTU *j* and *H_i_* is the total number of BLAST hits for spacer *i* across all categories. The ratio *h_ij_*/*H_i_* thereby dictates the fraction of a spacer hits mapping to a specific PTU. This reflects the mean proportion of targeting directed toward that specific PTU across the dataset.

## Author Contributions

AF-G, PG-F, IG-F, NB, DLG-H performed the experiments; DLG-H, RL, UK, ML supervised laboratory work; AF-G, ML, DB wrote MS; ML, DB conceptual design.

## Acknowledgments

Work in ML lab was supported by grant PID2023-147192OB-I00 from MICIU/AEI/10.13039/501100011033/ and ERDF/EU. DB was supported by European Research Council (no. 101044479) and Agence Nationale de la Recherche (no. ANR-10-LABX-62-IBEID). UK was supported by the JPIAMR TEXAS and the JPIAMR SEARCHER project funded by the German Bundesministerium für Forschung, Technologie & Raumfahrt under grant numbers 01KI2401 & 01KI2404A, as well as the ONE- BRIDGE project funded by the EU4Health Program (EU4H) under project number 101233407. Responsibility for the information and views expressed in the manuscript lies entirely with the authors. AF-G and PG-F were recipients of predoctoral appointments from the University of Cantabria. We are grateful to Harris Wang for sharing plasmids and unpublished information. We thank Olatz Irastorza for technical support, Eva Ceballos from SONINGEO for kindly providing wastewater samples, and the laboratory of M.P. Garcillán-Barcia for providing the database of plasmids with PTUs.

