## Supplementary information for "Tracking the trajectory of individual conjugative plasmids in bacterial communities using CRISPR acquisition-based recording"

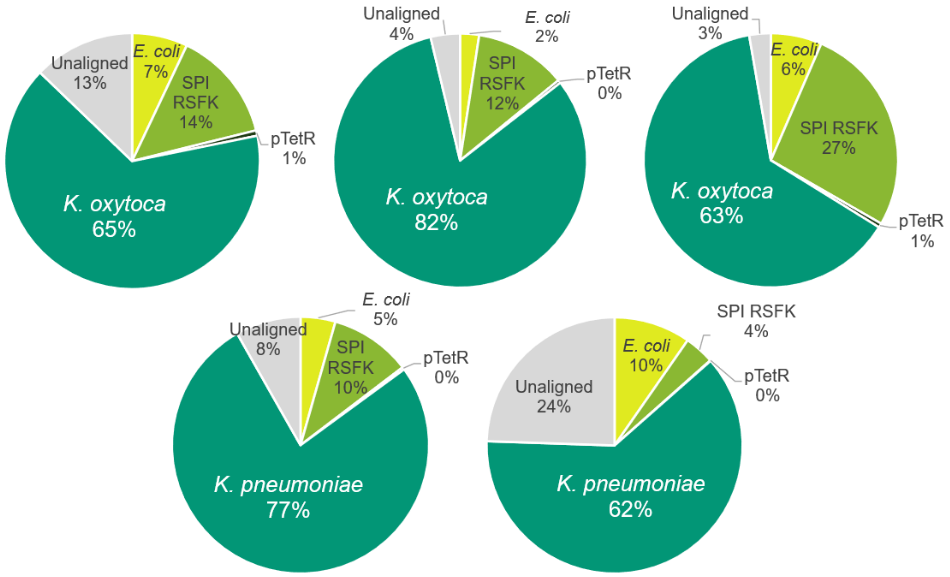


**Supplementary Figure 1. Spacer acquisition profiles from independent conjugations of SPI RSFK into *Klebsiella* recipients.** Pie charts display the relative proportions of acquired spacers mapping to different genetic sources across individual biological replicates. SPI RSFK was mobilized from an *E. coli* donor into *K. oxytoca* (top row, three independent experiments) and *K. pneumoniae* (bottom row, two independent experiments). Slices indicate the percentage of spacers targeting the recipient chromosome (*K. oxytoca* or *K. pneumoniae*), the *E. coli* donor chromosome, SPI RSFK plasmid, pTetR helper plasmid, or unaligned sequences.

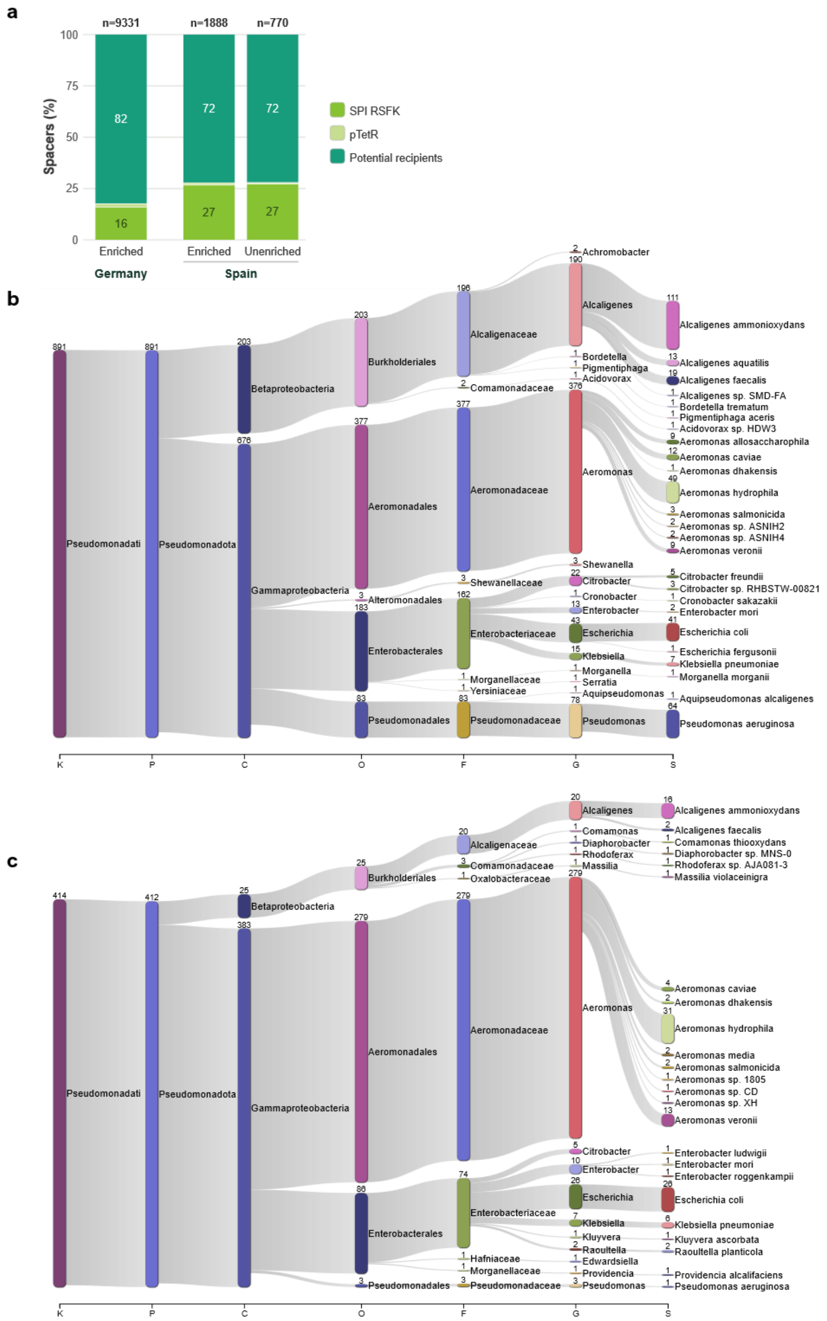


**Supplementary Figure 2. Origin and taxonomic classification of community-derived spacers acquired by SPI RSFK. a** Proportion of recovered spacers mapping to the donor plasmids (SPI RSFK and pTetR) and those potentially originating from the wastewater community in three conjugations, which correspond to matings with two wastewater communities: one from Germany (Dresden), under LB enrichment, and one from Spain (Lloredo), under both LB-enriched and unenriched conditions. The total number of spacers (n) is indicated above each bar **b, c** Sankey diagrams illustrating the taxonomic profiles of spacers acquired following the conjugation of SPI RSFK into the wastewater community from Spain. The assays were performed in parallel using either the LB-enriched microbial community (b) or the raw, unenriched concentrated wastewater (c). The diagrams span taxonomic ranks from Kingdom (K) down to species (S), displaying the 25 most abundant taxa per rank. Taxonomic assignments were performed using an LCA algorithm applied to BLAST hits sharing the highest bitscore.

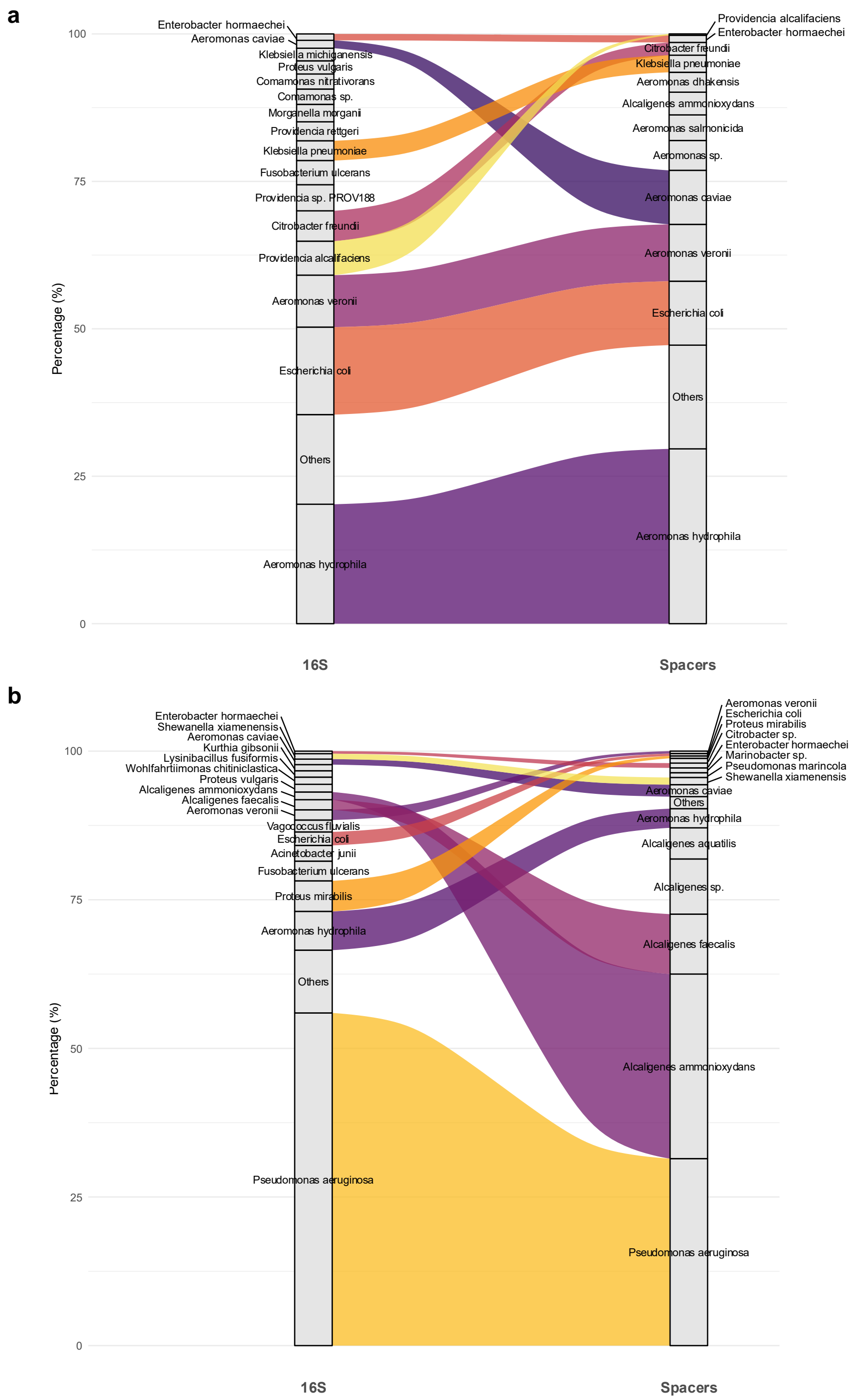


**Supplementary Figure 3. Correlation between microbial community composition and spacer acquisition by SPI RSFK in conjugations to the wastewater sample from Spain.** Alluvial diagrams comparing the relative abundance of bacterial species in the wastewater community, determined by 16S rRNA sequencing (left), with the frequency of acquired spacers assigned to those species via Best BLAST Hit (right). Matings were performed in parallel using either the LB-enriched microbial community (a) or the unenriched, concentrated raw wastewater (b). Flow widths are proportional to the relative frequency of each taxon across both datasets. Species outside the top 20 most abundant are grouped as ‘Others’.

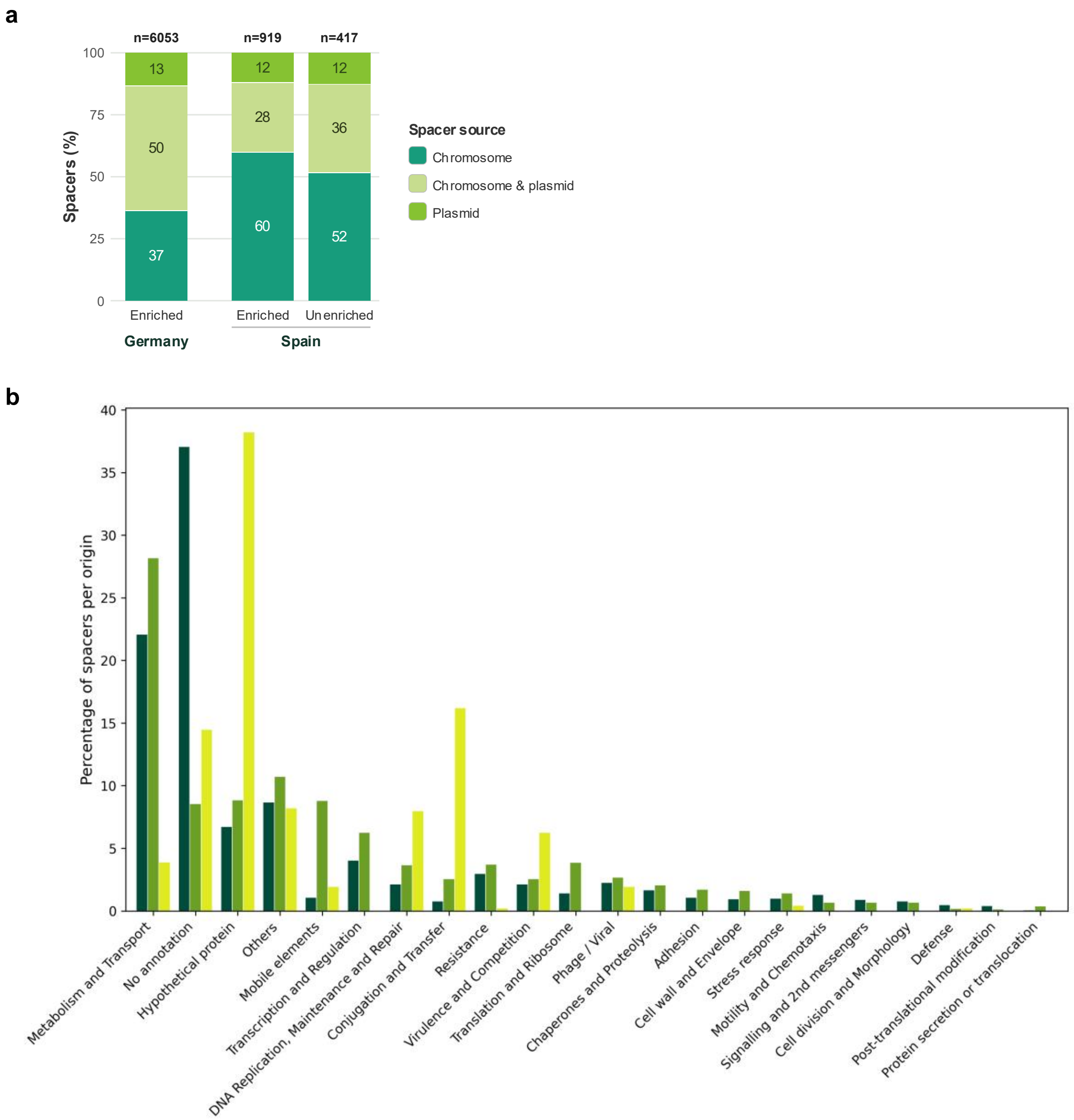


**Supplementary Figure 4. Genomic origin of the acquired spacers and functional categorization of the genes they target, for SPI RSFK conjugated into wastewater communities. a** Stacked bar chart of the inferred genomic origin of the acquired spacers in the three wastewater matings. For each sample, bars show the percentage of spacers matching exclusively chromosomal sequences, exclusively plasmid sequences, or both. The total number of spacers (n) is indicated above each bar. The Germany sample combines four independent sequencing runs, the LB-enriched Spain sample two, and the unenriched Spain sample a single run. **b** Functional annotation of the genomic regions targeted by the spacers from the Germany sample. Spacer targets are grouped by their inferred genetic origin, and bars show, within each origin category, the percentage of spacers aligning to genes assigned to each functional class.

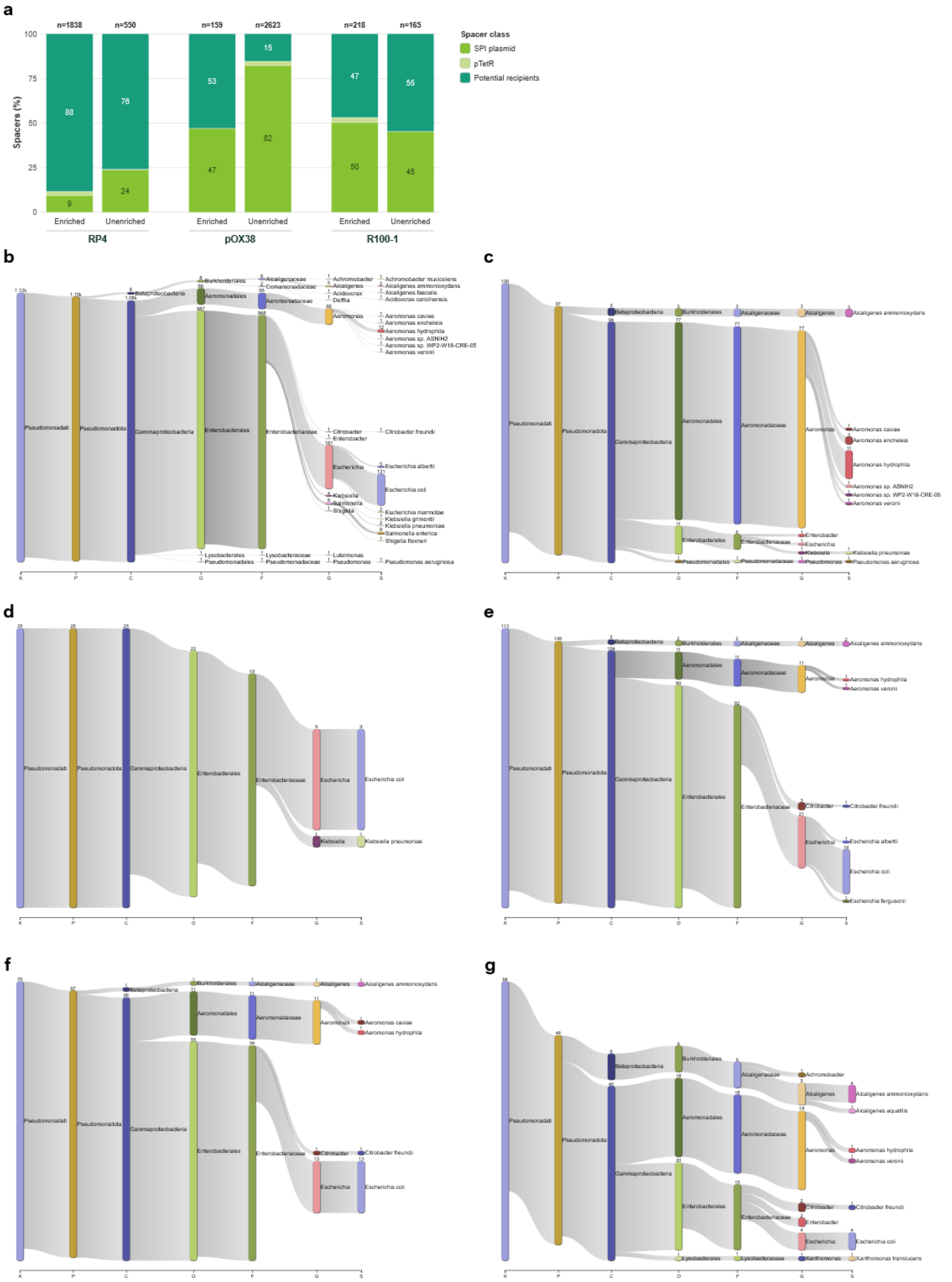


**Supplementary Figure 5. Origin and taxonomic profiling of the spacers acquired by the three conjugative SPI plasmids in wastewater.** SPI RP4, SPI pOX38 and SPI R100-1 were conjugated into the second wastewater community (Spain) under LB-enriched and unenriched conditions. **a** Proportion of recovered spacers mapping to the cognate SPI plasmid, to the pTetR helper, or remaining unassigned to either and therefore taken forward as potential recipient-derived spacers. **b-g** Sankey diagrams illustrating the taxonomic classification of the community-derived spacers, for SPI RP4 (b, c), SPI pOX38 (d, e) and SPI R100-1 (f, g). Assays were conducted in parallel using either the LB-enriched microbial community (left column: b, d, f) or the unenriched, concentrated raw wastewater (right column: c, e, g). The diagrams trace taxonomic assignments from Kingdom (K) down to species (S), performed using an LCA algorithm applied to BLAST hits sharing the highest bitscore. The enriched RP4 and unenriched pOX38 datasets each combine two independent sequencing runs of the same sample, whereas all other datasets derive from a single run.

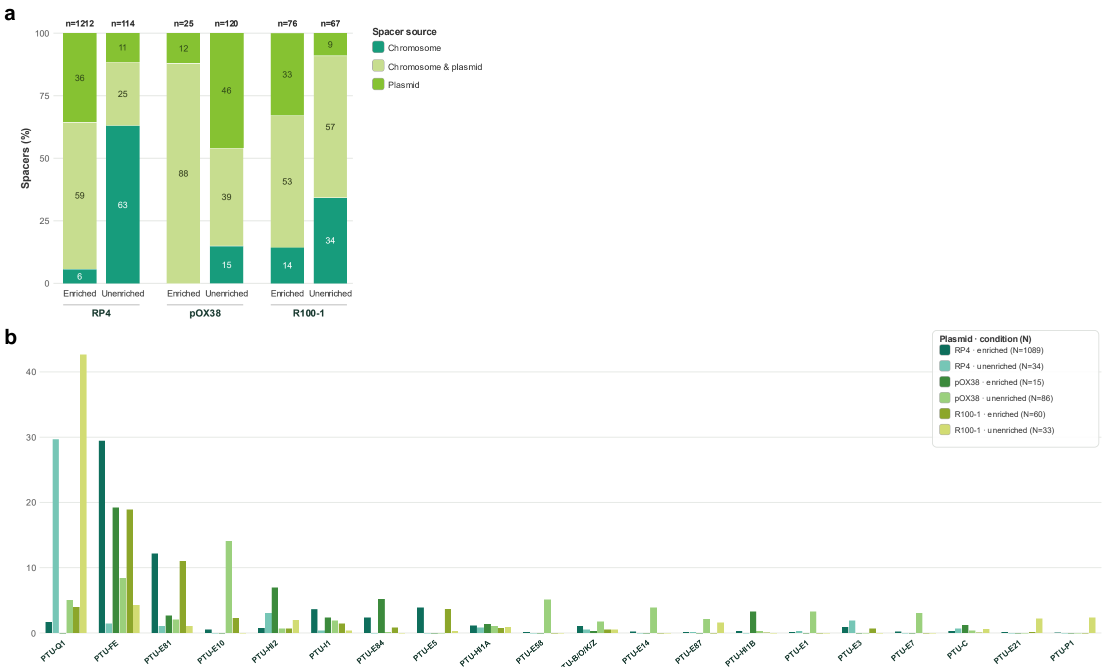


**Supplementary Figure 6. Genomic origin and PTU classification of plasmid-derived spacers acquired by the three conjugative SPI plasmids in wastewater.** The three conjugative SPIs were mated with the Spain wastewater community under both LB-enriched and unenriched conditions. **a** Inferred genomic origin of the acquired spacers, shown as the percentage matching exclusively chromosomal sequences, exclusively plasmid sequences, or both. Total numbers of spacers (n) are shown above the bars. **b** Weighted presence of the 20 most abundant PTUs among the plasmid-derived spacers across the six conjugations (parallel bars); candidate and provisional PTUs were excluded from the plot, and the number of spacers with at least one PTU-assigned hit (N) is indicated per conjugation in the legend. The enriched RP4 and unenriched pOX38 datasets each combine two independent sequencing runs of the same sample, whereas all other datasets derive from a single run.

**Supplementary Table 1. Plasmids constructed in this work.**

| Descriptive name[*1] | Laboratory Name | Description | Construction method[*2] | Vector source[*3] | Insert/s or excised fragment | Insert/s source |
|---|---|---|---|---|---|---|
| **pAF38** | pAF38 | pRec-ΔLacI::Minimal CRISPR array; $Cm^R$ | GA | pRec-ΔLacI | CRISPR array | Primer overhangs |
| **pAF48** | pAF48 | pAF38 Δ*tetR*::*aacC1*; $Cm^R$ $Gm^R$ | GA | pAF38 (without *tetR*) | *aacC1* | pBBR6 |
| **pTetR-Ap** | pAF51 | pUC18::*tetR* | GA | pUC18 | *tetR* with its promoter | pRec-ΔLacI |
| **pTetR-Cm** | pMTX1002 | pRec-ΔLacI Δ*cas1-2*; $Cm^R$ | RC | pRec-ΔLacI | pRec digested with ZraI and self-ligated to interrupt and partially remove *cas1-2* | |
| **SPI pOX38** | pGUR33 | pOX38 Δ*finO*::P*tet-cas1-2*:: *aacC1*::CRISPR array; $Cm^R$ $Gm^R$ | DWR | pOX38 | PCR cassette including adaptation module | pAF48 |
| **SPI R100-1** | pGUR35 | R100-1 Δ*tetA-R*::P*tet-cas1-2*:: *aacC1*::CRISPR array; $Cm^R$ $Gm^R$ | DWR | R100-1 | PCR cassette including adaptation module | pAF48 |
| **SPI RP4** | pAF55 | RP4 Δ*tetA-R*::P*tet-cas1-2*::*aacC1*::CRISPR Array; $Ap^R$ $Km^R$ $Gm^R$ | DWR | RP4 | PCR cassette including adaptation module | pAF48 |
| **SPI RSFK** | pMTX1001 | SPI RSFK-TetR Δ*tetR*; $Km^R$ | RC | pRec-ΔLacI | pAF41 digested with PstI and self-ligated to remove *tetR* | |
| **SPI RSFK oriT-** | pAF57 | SPI RSFK *oriT*$^-$; $Km^R$ | GA | SPI RSFK | Mutations in *nic* site of *oriT* | Primer overhangs |
| **SPI RSFK-TetR** | pAF41 | RSFK::P*tet-cas1-2*::*tetR*::CRISPR array; $Km^R$ | GA | RSFK | - P*tet-cas1-2*::*tetR*<br>- Minimal CRISPR array | - pRec-ΔLacI<br>- Primer overhangs |

*1 Commonly used plasmids have been assigned a descriptive name to facilitate readability.

*2 GA: Gibson Assembly; RC: Restriction Cloning; DWR: Datsenko and Wanner Recombination.

*3 All vectors for Gibson assembly were PCR amplified.

**Supplementary Table 2. Oligonucleotides used in this work.**

| Name | Oligonucleotide sequence (5' to 3') |
|---|---|
| **Oligonucleotides used in plasmid construction*[1]** | |
| **Construction of pAF38** | |
| **pRec_F_2.0** | **CCAGATATCGACGTCTAAGAAAC** |
| **pRec_vector_R** | **GATACGCCTATTTTTATAGGTTAATGTCATG** |
| **pRec_array_F** | AAAATGCTTTAAGAACAAATGTATACTTTTAGAGAGTTCCCCGCGCCAGCGGGGATAAACCG**CAGCAAAAGGCCAGGAACC** |
| **pRec_array_R** | ACTCTCTAAAAGTATACATTTGTTCTTAAAGCATTTTTCCCATAAAAACAACCCACCAACCTT**TCACATGTTCTTTCCTGCGTTATC** |
| **Construction of SPI RSFK-TetR** | |
| **pRec_RSFK_F** | CAATGTAACATCAGAGATTTTGAGACACAACGTGGCTTTG**CCAGATATCGACGTCTAAGAAAC** |
| **pRec_RSFK_R** | ATGTATGCTCTTCTGCTCCTGCAGCTAATGGATCACCGCA**TCACATGTTCTTTCCTGCGTTATC** |
| **array_RSFK_F** | AATGCTTTAAGAACAAATGTATACTTTTAGAGAGTTCCCCGCGCCAGCGGGGATAAACCG**CTGGAGCATGGCTTTCTACG** |
| **array_RSFK_R** | CTAAAAGTATACATTTGTTCTTAAAGCATTTTTCCCATAAAAACAACCCACCAACCTT**CCGCCCGCATTGGAGAAATTC** |
| **RSFK_F** | **TGCGGTGATCCATTAGCTGC** |
| **RSFK_R** | **CAAAGCCACGTTGTGTCTC** |
| **Construction of SPI RSFK oriT-** | |
| **Delta_nic_Fw** | **CGAAGAGAAACCGGTAAG**CTT**G**T**CCTCCCCTACAAAGTAGGGTCG** |
| **Delta_nic_Rv** | **CGACCCTACTTTGTAGGGGAGG**A**C**AAG**CTTACCGGTTTCTCTTCG** |
| **Cas1-2_R** | **TCAGCTACTCCGATGGCCTGC** |
| **array_genoma_F** | **TCCTGTTTCACTGGGAGATGCAG** |
| **Construction of pAF48** | |
| **rrnB_F** | **CAAATAAAACGAAAGGCTCAGTC** |
| **T7Te_R** | **CGCAGAAAGGCCCACCC** |
| **Gm_F** | TAGAGTCACACTGGCTCACCTTCGGGTGGGCCTTTCTGCG**TCGCGGCCATCGTCCAC** |
| **Gm_R** | GAAAGGCCCAGTCTTTCGACTGAGCCTTTCGTTTTATTTG**TTAGGTGGCGGTACTTGGG** |
| **Construction of SPI RP4** | |
| **RP4_Wanner_F** | CTACATGGCTCTGCTGTAGTGAGTGGGTTGCGCTCCGGCAGCGGTCCTGA**AATAGGCGTATCACGAGGC** |
| **RP4_Wanner_R** | TTGCCGGAAGTCGCCTTGACCCGCATGGCATAGGCCTATCGTTTCCACGA**GCCAGCAACGCGGCC** |
| **Construction of SPI pOX38** | |
| **pOX38_Wanner_F** | GTGTGGTATCTTTGGCCGCAAGAGGATGATTCGTCAGAGGCAGAACACAC**CAGATATCGACGTCTAAGAAAC** |
| **pOX38_Wanner_R** | GTAAGATCAGCTTTCTGAACGGGTGATGGCCTTCAGCGCCCTGCGCAGT**GTTCCTGGCCTTTTGCTG** |
| **Construction of SPI R100-1** | |
| **R100_Wanner_F** | CATCACGGAAAAAGGTTATGCTGCTTTTAAGACCCACTTTCACATTTAAGC**CAGATATCGACGTCTAAGAAAC** |

| | |
|---|---|
| **R100_Wanner_R** | CAAGAGGGTCATTATATTTCGCGGAATAACATCATTTGGTGACGAAATAA**GTTCCTGGCCTTTTGCTG** |
| **Construction of pTetR-Ap** | |
| **pUC18_F** | **ACTGGCCGTCGTTTTACAAC** |
| **pUC18_R** | **GGCGCTCTTCCGCTTCC** |
| **TetR_F** | GAGCGCAGCGAGTCAGTGAGCGAGGAAGCGGAAGAGCGCC**TTTATAGGTACCCGTGGATCCTCTAG** |
| **TetR_R** | GGTTTTCCCAGTCACGACGTTGTAAAACGACGGCCAGT**ATTTGTCCTACTCAGGAGAGCG** |
| **Oligonucleotides used for the detection of array expansion*[2]** | |
| **CRISPR array of plasmids SPI RSFK-TetR and SPI RSFK** | |
| **RSFK_seq_F** | **GGGAAAGGAATTGCCGGGC** |
| **RSFK_seq_R** | **CTCGGGCAGAAACCGCGACGC** |
| **CRISPR array of plasmid SPI RP4** | |
| **pRec_seq** | **TATCAGCTCACTCAAAGGCGG** |
| **RP4_compr_R** | **CCAACCGCACTCCTAGGGC** |
| **CRISPR array of plasmid SPI pOX38** | |
| **pRec_seq** | **TATCAGCTCACTCAAAGGCGG** |
| **pOX38_R2** | **CTGCGCAGTGTTCCTGG** |
| **CRISPR array of plasmid SPI R100-1** | |
| **pRec_seq** | **TATCAGCTCACTCAAAGGCGG** |
| **R100_1_Rv** | **TCGCGGAATAACATCATTTGG** |
| **Universal reverse primer for nested PCRs** | |
| **RepeatDegen_R** | **CCGCTGGCGCGGGGAACTCV** |
| **RepeatDegen_Tail_R** | ATAGTCA**TGGCGCGGGGAACTCV** |
| **Oligonucleotides used for 16S rRNA amplification** | |
| **1492R** | **TACGGYTACCTTGTTACGACTT** |
| **27F** | **AGAGTTTGATYMTGGCTCAG** |

*[1] For oligonucleotides used in plasmid construction, the region that anneals with the template is shown in bold, while the primer tails are indicated in regular text.

*[2] In nested PCRs with degenerate oligonucleotides for the selective amplification of expanded arrays, PCR2 is done in each case using the forward oligonucleotide specific to that array and the oligo RepeatDegen_R or RepeatDegen_Tail_R (with overhang to prevent formation of chimeric arrays).
